# Parietal cortex is causally required for state-dependent decisions

**DOI:** 10.1101/2024.05.23.595581

**Authors:** Akhil C. Bandi, John P. McCann, Rachel M. Carpenter, Mason J. Lower, Jordyn N. Long, Caroline A. Runyan

## Abstract

During perceptual decision-making tasks, behavioral performance and its neural correlates vary with changes in internal states such as arousal, motivation, and strategy. It is not yet clear whether these state fluctuations differently impact the distributed neural processes that underlie task performance, from sensory processing to control of motor outputs. The posterior parietal cortex (PPC) is an association-level cortical region implicated in sensorimotor transformations, but evidence of its causal requirement in decision-making tasks has been contradictory. We trained mice to perform a navigation-based sound localization task and asked how natural fluctuations in behavioral state related to neural processing in association areas during decision-making. Behavioral performance in expert-level mice was not static but instead transitioned between periods of near-perfect performance and biased, less accurate performance. Using a hidden Markov model, we could reliably define these as distinct strategies that included a high-performance state where mice used relevant stimulus information to inform choices, and two biased states where mice weighted stimulus information less strongly. Optogenetic inactivation of PPC decreased task performance accuracy, and the model captured the resulting change in behavioral strategy as a reduction in the weighting of the auditory cues and an increase in behavioral bias, in predicting mice’s decisions. Two-photon calcium imaging revealed that performance states strongly influenced population activity patterns in PPC, but not primary auditory cortex (AC). Surprisingly, activity of individual PPC neurons was better explained by external inputs and behavioral variables during biased behavioral performance, while shared variability across neurons in PPC was strongest in the high-performance state. Together, these findings suggest that neural activity in parietal cortex is causally required for decisions and is linked to behavioral performance states.

## Introduction

Animals do not maintain high levels of alertness and behavioral performance consistently for long periods of time, but instead fluctuate in arousal and motivation levels, which affects perceptual and decision-making performance^1–3^. Shifting performance during decision-making tasks can be modeled as discrete states with alternate strategies guiding choice behavior^4,5^. Animals capable of expert-level task performance predominantly occupy a high-performance state, using relevant sensory information to guide behavioral choices. Yet even these expert animals can occupy suboptimal performance states when satiated or less motivated, responding with behavioral bias instead of using relevant stimulus information. These periods of suboptimal performance are characterized by increased variability in correlates of arousal such as pupil diameter and uninstructed movements^6,7^.

The posterior parietal cortex (PPC) is an association-level area that integrates multimodal sensory information^8–10^ and encodes upcoming behavioral choices^11–24^. While PPC has long been proposed as a key node the brain’s decision network^8,25,26^, studies testing its causal role in perceptual decision-making have yielded inconsistent findings^27–32^. Some studies suggest PPC activity to be entirely dispensable for perceptual decision-making task performance^27,33^, while others demonstrate its necessity for both the evaluation of sensory information and its transformation to motor output^28–30^. We hypothesize that neural activity states in PPC and latent behavioral performance states, such as behavioral strategy, are causally linked. The goal of our study is to test the effects of PPC inactivation on decision-making strategy, and to determine the effects of these behavioral states on neural encoding in PPC.

We trained mice to perform a navigation-based decision-making task using auditory cues, finding that even well-trained mice shifted between behavioral performance states with high accuracy or more biased choices. Optogenetic inactivation of PPC shifted performance toward biased strategies where mice were less likely to use the relevant sound cue to guide their choices. Population activity patterns differed significantly across performance states in PPC in two key ways: 1) PPC activity could be used to decode the latent behavioral state, and 2) shared variability across neurons was greater in PPC during the high-performance state. In contrast, population activity in primary auditory cortex (AC) was unaffected by latent performance state. Together, our results reveal that activity in PPC is causally linked to performance strategies used by mice in this auditory decision-making task.

## Results

### Causal requirement for parietal cortex in an auditory decision-making task

Mice performed a virtual reality (VR) based T-maze task, using the locations of sound stimuli to guide left-right choices^24^. As mice ran down the virtual corridor a sound cue was played from one of eight possible locations. Mice reported whether the sound came from a left or right direction by turning in that direction at the T-intersection (Figure 1A). Expert mice learned to accurately categorize the locations of the sound cues, making greater than 70% correct choices in a session, and used a subjective category boundary (as shown by the peak of the psychometric slope function) that closely matched the experimentally defined category boundary (N=5 mice, 69 total behavioral sessions; Figure 1B). Although animals performed well across an entire session, they did not perform at a constant level of behavioral accuracy in the task but instead fluctuated between periods of low task performance with increased choice bias and periods of high accuracy (Figure 1C).

**Figure 1.**
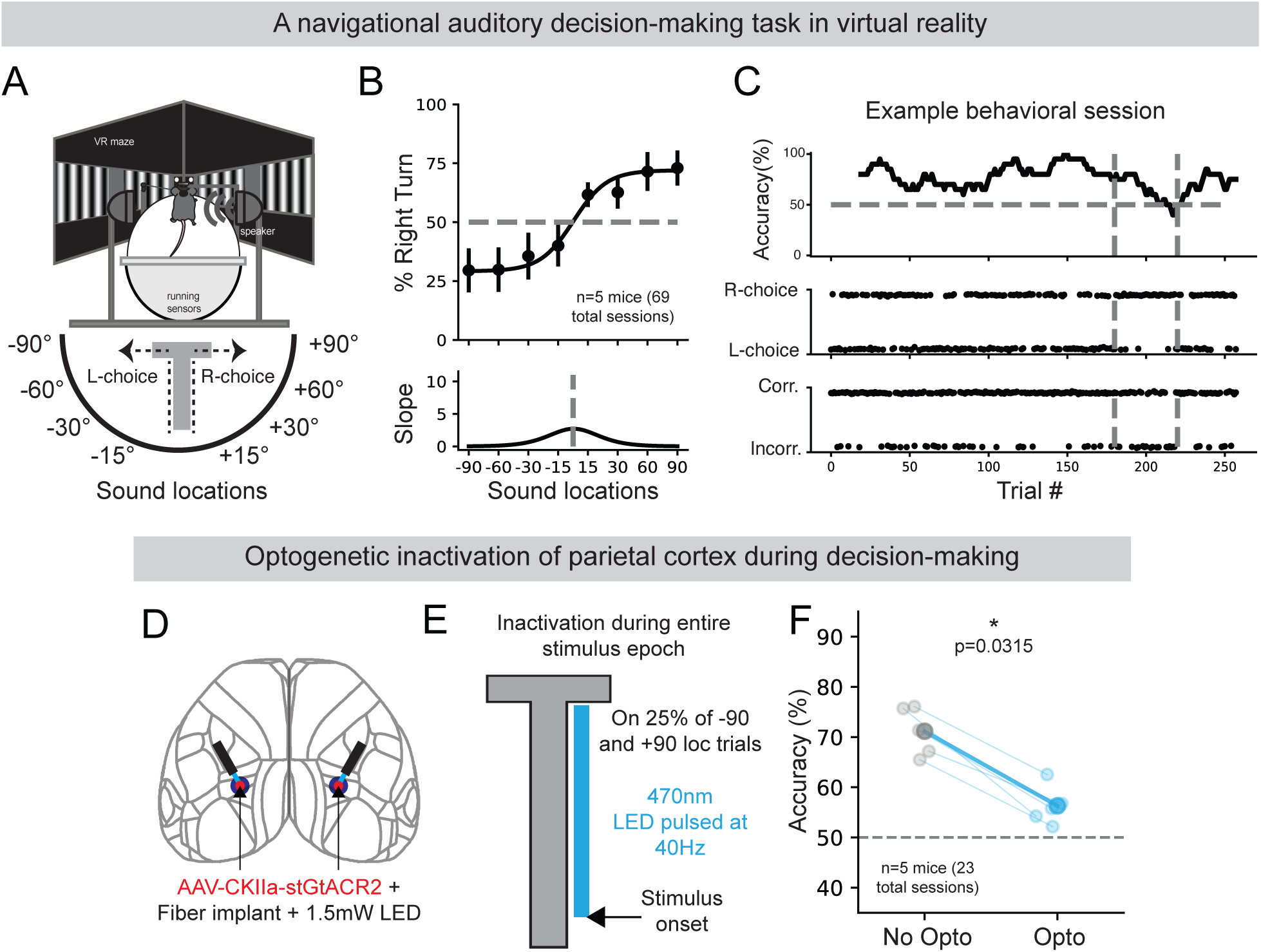
Parietal cortex was causally required for performance of an auditory decision-making task. **(A)** Mice were trained to report the location of a sound stimulus by turning in the direction of the sound in a virtual-reality T-maze. **(B)** Mice learned to categorize sound locations and performed the task (n=5 mice, 69 total sessions, mean ± s.e.m) as evidenced by the psychometric curve for the probability of a right-ward choice given a sound stimulus for a specific location. **(C, top)** Performance of an example mouse in one behavioral session presented as a 20-trial moving window of correct choice percentage. Vertical dashed lines indicate a period of trials where the mouse’s performance was low due to increased right-choice bias. **(C, middle)** Left or right choice identity for each trial. **(C, bottom)** Correct or incorrect identi-ty for each trial. **(D)** Schematic showing the location of bilateral AAV-CKIIa-stGtACR2 injections and optical fiber implants in PPC. **(E)** Schematic showing activation of stGtACR2 using 470nm to inactivate PPC activity from stimulus onset to the stem of the t-maze on 25% of −90 and +90 sound location trials. **(F)** Effect of PPC inactivation on task accuracy (n=5 mice, 23 total sessions, mean ± s.e.m, No-Opto mean = 71.2 ± 2.1, Opto mean = 56.3 ± 1.7, p= 0.0315, Wilcoxon signed rank test).

Next, we optogenetically inactivated PPC, to measure its requirement for accurate performance in the task. We injected a viral construct to express a soma-targeted anion-conducting channelrhodopsin in excitatory neurons (AAV1-CKIIa-stGtACR2^34^, Supplementary figure 1) and implanted optic fibers bilaterally into PPC (Figure 1D). This approach allowed us to use 470nm LEDs at 1.5mW to inactivate excitatory neuron activity in PPC. During task performance, photoinhibition was applied on 25% of −90 and +90 auditory location trials. Photoinhibition started at stimulus onset and stopped at the end of the stem of the T-maze (Figure 1E). Inactivating PPC reduced performance in the task (No-opto mean = 71.2±2.1%; Opto mean = 56.3±1.7%; p = 0.0315, Wilcoxon signed-rank test; N=5 mice, 23 total sessions; Figure 1F). When identical illumination patterns were applied to control animals not injected with stGtACR2, task performance was not impacted (Control No-opto mean = 71.6±9.3%; Control Opto mean = 73.2±8.1%; p = 0.48, Wilcoxon signed-rank test; N=3 mice, 15 total sessions; Supplementary figure 2).

In separate sessions, PPC was inactivated at different task epochs (first half of the maze, second half of the maze, or for 2 seconds during the inter-trial interval). Inactivation during the second half of the maze yielded the largest decrease in task performance (Supplementary figure 3). Furthermore, the effects of photoinhibition on task performance persisted for multiple sessions indicating the continued causal requirement of parietal function well after mice have acquired the task (N=3 mice, 15 total sessions; Supplementary figure 4). Taken together, these results show that parietal cortex activity is causally required for accurate performance of the task.

### Mice fluctuated between latent strategy states during the auditory decision-making task

To characterize the fluctuations in task performance that we observed (Figure 1C), and to identify potential changes in hidden states guiding behavioral performance, we used hidden Markov models with generalized linear model observations (GLM-HMMs^4^). We modeled the mouse’s decision-making strategy using a GLM-HMM with four inputs (Figure 2A): (1) the left-right location category of the auditory stimulus on a given trial; (2) the binary (left-right) choice made by the mouse on the previous trial; (3) whether the previous trial was correct or incorrect; and (4) a constant offset or choice bias. We chose the GLM-HMM approach because it allows us to simultaneously model how multiple behavioral factors influence decision-making while also capturing temporal dynamics through latent behavioral states that can switch over time, providing a more comprehensive framework than simpler models that either ignore state transitions or treat behavioral factors independently. Each state in the HMM corresponds to a Bernoulli GLM for a rightward choice given the four input predictors. We fit each GLM-HMM with varying numbers of latent states to choice behavior using 69 total behavioral sessions across 5 mice performing the eight-location VR sound localization task. The three-state GLM-HMM explained the data better than a model without latent states, as evidenced by higher log-likelihood and predictive accuracy on held-out test data. We limited the model to three states, because model performance increases plateaued with additional states (Supplementary figure 5A-B).

**Figure 2.**
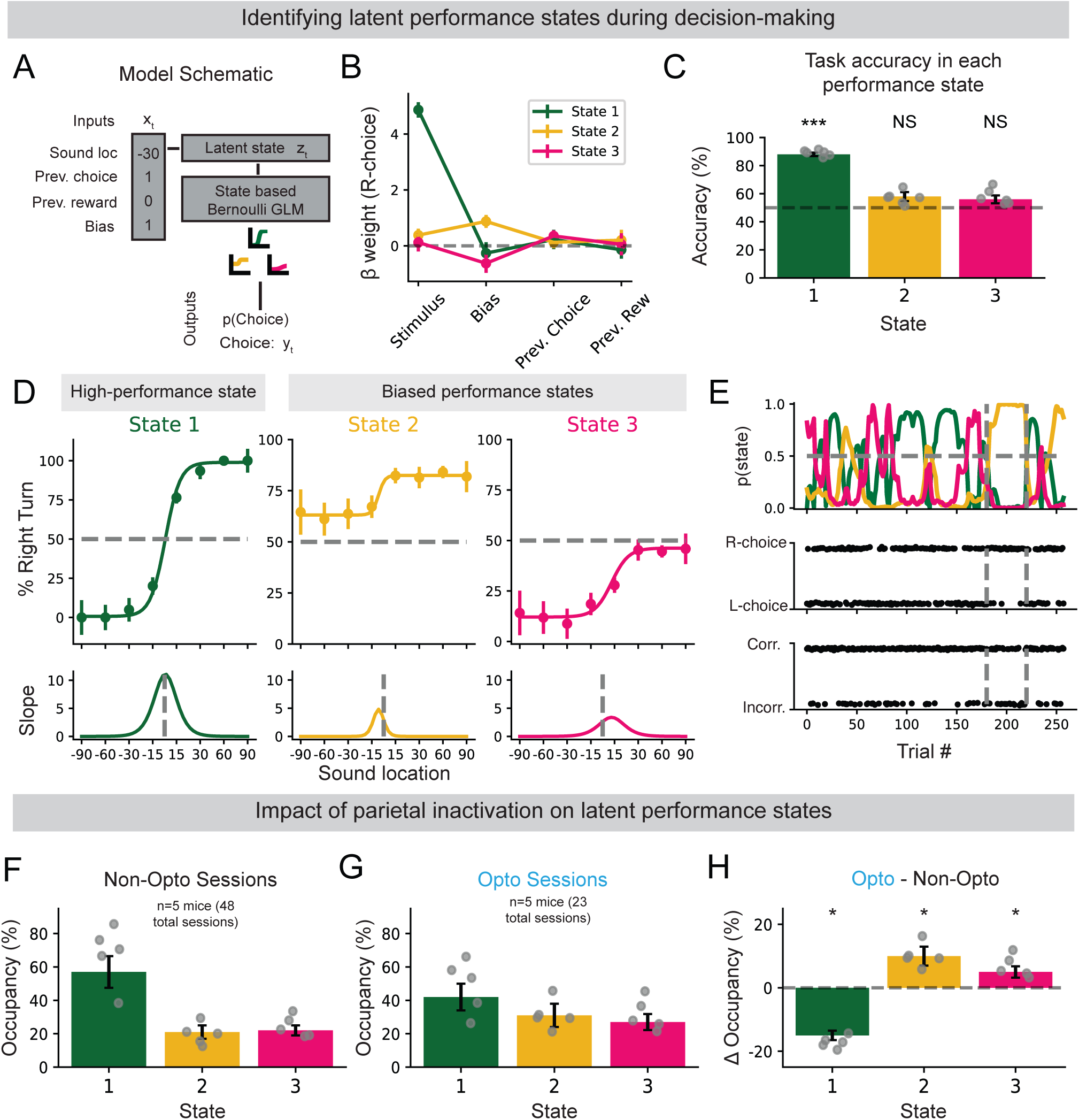
Behavioral deficits caused by parietal cortex inactivation impacted the occupation of latent performance states during decision-making. **(A)** Three-state GLM-HMM model with four input variables and three different GLMs corresponding to different decision-making strategies. **(B)** Inferred GLM weights for the three-state GLM-HMM. State 1 had a high weight for stimulus information, and states 2 and 3 had high and opposing weights for bias (n=5 mice, mean ± s.e.m). **(C)** Task accuracy across each GLM-HMM state (n=17,602 trials, mean ± s.e.m; state 1 = 88.0 ± 1.2, p=1.0 x 10^-5^; state 2 = 58.0 ± 6.2, p=0.054; state 3 = 56.0 ± 2.2, p=0.089; One-sample Wilcoxon signed rank test). **(D)** Per-state psychometric curves for the probability of a right-ward choice given a sound stimulus for a specific location. The differ-ent psychometrics across the three states highlight differences in decision-making strategy and performance. **(E)** Posterior state probability for the example session shown in figure 1C. The three-state model successfully identified the period of poor task performance and classified the animal’s behavior as being in state 2, which was characterized by rightward bias. **(F)** Fractional occupancies for each state across all non-inactivation behavioral sessions (n=5 mice, mean ± s.e.m, state 1 = 57.1 ± 10.8, state 2 = 20.7 ± 4.4, state 3 = 22.7 ± 3.8). **(G)** Fractional occupancies for each state across all behavioral trials from inactiviation sessions (n=5 mice, mean ± s.e.m, state 1 = 41.9 ± 8.6, state 2 = 31.3 ± 7.6, state 3 = 37.2 ± 2.8). **(H)** Change in state occupancy for each mouse (occupancy on inactivation sessions - occupancy on non-inactivation sessions). Decrease in state 1 occupation and increase in state 2 and 3 occupation (n=5 mice, Δ mean ± s.e.m, state 1 = −16.7 ± 4.7, p = 0.010; state 2 = 9.1 ± 5.2, p = 0.031; state 3 = 6.1 ± 2.8, p = 0.040; One-sample Wilcoxon signed rank test).

The three GLM-HMM states defined three distinct behavioral strategies where mice either used sound stimuli to more accurately perform the task (state 1) or instead responded with left or right choice bias and lower performance accuracy (states 2 and 3). This was evident in the inferred GLM weights related to the mouse’s choice in each state. In state 1, the sound stimulus location input was strongly weighted, while previous choice, previous reward outcome, and choice bias had negligible weights. In contrast, the GLMs for state 2 and state 3 had smaller magnitude weights for stimulus location, but higher magnitude weights for the left-right bias input (Figure 2B). When mice occupied state 1, task performance was high (88.0±1.2% correct; p=1.0 x 10^-5^, One-sample Wilcoxon signed rank test, Figure 2C), and when mice occupied states 2 and 3, task performance was lower (state 2 mean = 58±6.2% correct, p=0.054; state 3 mean = 56±2.2% correct, p=0.089; One-sample Wilcoxon signed rank test; Figure 2C).

After identifying behavioral states, we examined choice formation as measured through running kinematics and a convolutional neural network trained to predict directional choices from movement trajectories^20^. While running velocities in the x-axis remained consistent across states (Supplementary figure 6A), y-axis velocity was significantly reduced in the high-performance state 1 compared to the biased performance states (Supplementary figure 6B, p=0.015 and p=0.026, respectively; Mann-Whitney U-Test). To quantify choice formation timing, we trained a convolutional neural network to predict upcoming left/right choices using kinematic variables throughout each trial (Supplementary figure 6). The latency to dynamic choice, defined as when the model’s prediction confidence exceeded threshold (0.9 for left, 0.1 for right), was significantly longer in state 1 compared to both state 2 (p=0.024) and state 3 (p=0.011), with no difference between the suboptimal states (Supplementary figure 6D). This indicates that during high-performance periods, mice took longer to commit to a directional decision.

We further assessed the decision-making strategies associated with each state by plotting psychometric curves, relating the probability of a rightward choice to the stimulus location (Figure 2D). The state 1 psychometric had a steeper slope at the left-right category boundary, indicating that the mouse more similarly categorized both easy and difficult sound locations. By comparison, the psychometric curves measured during states 2 and 3 had more shallow slopes and were shifted up or down, reflecting rightward and leftward choice biases, respectively. To characterize the temporal structure of how mice transitioned between states during performance of the task, we plotted the GLM-HMM state’s posterior probability for every trial in the behavioral session (Figure 2E). States persisted for many trials in a row (state 1: 20.2±8.2 trials, state 2: 8.6±2.0 trials, state 3: 6.9±1.3 trials), and multiple state transitions occurred throughout a behavioral session (11.3±6.9 state switches). In the majority of trials, the most predicted state had a high posterior probability (0.9 on 87% of trials), and so there were few trials with ‘mixed’ or competing state predictions. Overall, the three-state GLM-HMM revealed that mice trained to expertly perform the task transitioned between high-performance and biased-performance latent states of behavioral performance, characterized by unique strategies governing behavioral choices.

### PPC inactivation led to biased latent performance states during the task

Since PPC inactivation reduced performance accuracy in the task (Figure 1F), we next considered whether the GLM-HMM approach could reveal the impacts of PPC inactivation on behavioral strategy. In control sessions (when PPC activity was not manipulated), state 1 was the most frequently occupied state across all behavioral sessions (57.1±10.8%; Figure 2F), states 2 and 3 were less frequently occupied (20.7±4.4% and 22.7±3.8%; Figure 2F). We then fit the GLM-HMM to sessions in which PPC was optogenetically inactivated. State occupation was modulated by PPC inactivation, as state 1 occupation was reduced, and states 2 and 3 were increased (state 1: 41.9±8.6%, state 2: 31.3±7.6%, state 3: 37.2±2.8%; Figure 2G). Thus, when PPC was optogenetically inactivated, state occupation was significantly modulated (State 1Δ = −16.7±4.7%, p=0.01; state 2Δ = 9.1±5.2%, p=0.031; state 3Δ = 6.1±3.8%, p=0.040; One-sample Wilcoxon signed rank test, Figure 2H). When inactivation was restricted to particular task epochs, only inactivation during the second half of the maze altered latent state occupation, while inactivation earlier in the trial or during the inter-trial-interval had no impact (Supplemental figure 7). Together, these results suggest that parietal cortex activity is necessary for sustaining the high-performance behavioral state characterized by optimal stimulus weighting.

### PPC population activity patterns varied with performance state

Having found that disrupting activity in PPC altered the latent performance state during the task, we next tested whether a signature of the mouse’s current performance state exists in neural activity. We used in-vivo calcium imaging to monitor the activity of GCaMP6^+^ neurons in PPC, as well as in auditory cortex (AC) for comparison (Figure 3A). First, we tested if population activity patterns of AC and PPC neurons distinguished between the mouse performing the task in the high-performance (state 1) or biased performance state (states 2 or 3). We trained binary support vector machine (SVM) decoders to classify each trial as high-performance (defined as state 1 occupation) or biased (combining states 2 and 3, to balance choice direction) from AC or PPC neuronal population activity during that trial. We then calculated the decoding accuracy using five-fold cross validation (Figure 3B). Importantly, we equally balanced the number of left and right stimulus and left and right choice trials across engaged and biased states, to allow the decoder to classify behavioral strategy state independently of the mouse’s left-right (See methods section “Balancing trial conditions to decorrelate stimulus, choice, and behavioral performance state”). The accuracy of decoding of performance state using PPC activity was better than chance-level decoding accuracy (N=8 datasets, mean=76.14±9.13%, p= 4.1 x 10^-5^, One-sample Wilcoxon signed rank test, Figure 3C), while decoding of performance state using AC activity was similar to chance performance (N=7 datasets, mean=53.48±4.73%, p= 0.67, One-sample Wilcoxon signed rank test, Figure 3C). To summarize, population activity in PPC but not AC discriminated between the mouse’s behavioral performance state, meaning that population activity patterns in PPC were different when mice used sensory stimuli to inform choices (high-performance state, 1), as opposed to following the biased performance strategy (biased states, 2 and 3).

**Figure 3.**
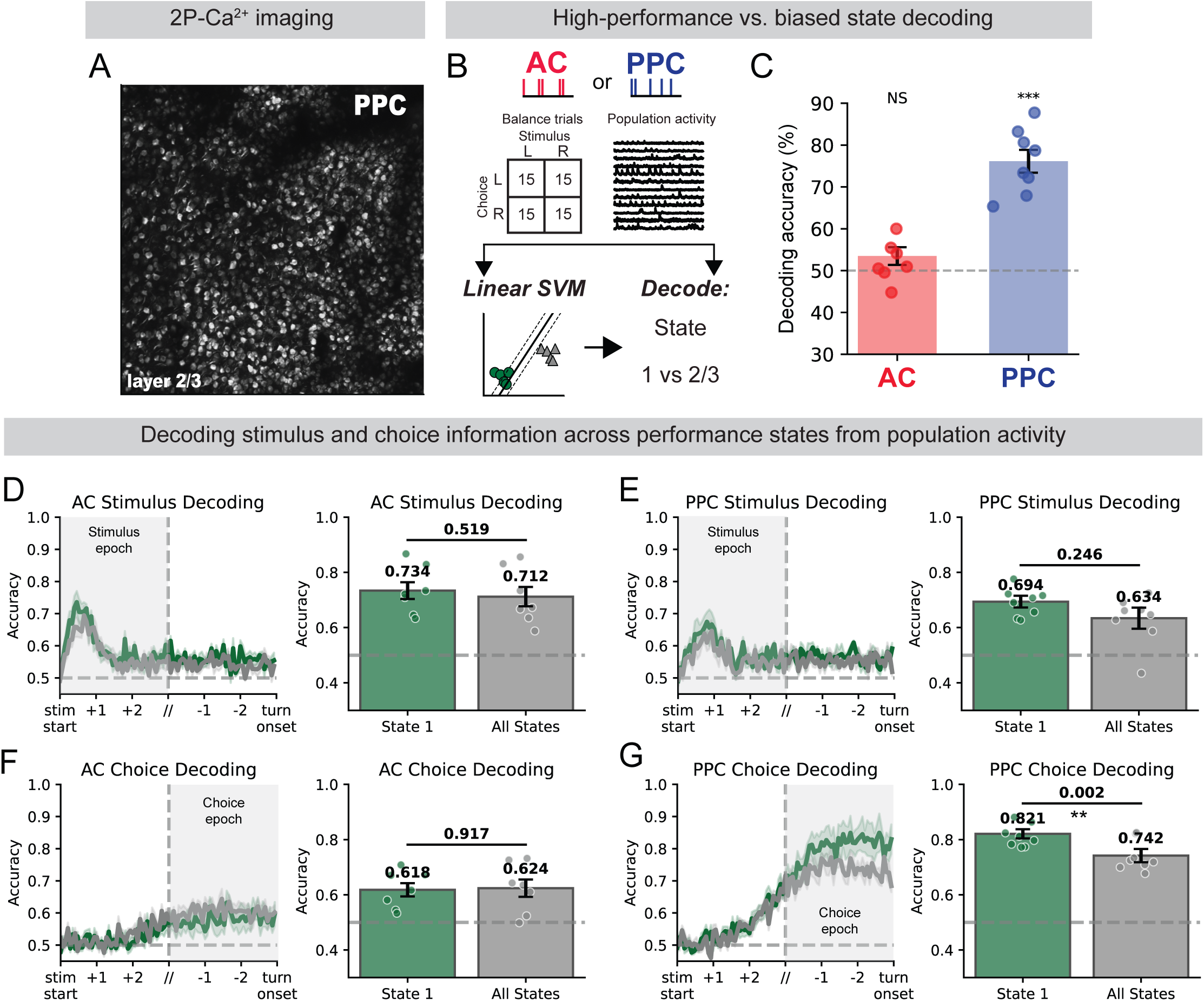
PPC population activity patterns varied with performance state. **(A)** Example field of view from 2P-Ca^2+^ imaging of PPC. **(B)** Schematic of using a linear SVM to decode performance state information from AC and PPC population activity. The number of left and right stimulus/choice trials were balanced across states. **(C)** SVM decoding accuracy of the classification of a trial as occuring during the high performance or biased states from PPC or AC population activity during that trial (n=7 recording sessions for AC and 8 for PPC; AC mean = 53.48 ± 4.73, p= 0.67; PPC-mean = 76.14 ± 9.13, p= 4.1 x 10^-5^, Wilcoxon signed rank test). **(D)** SVM decoding accuracy of stimulus information from AC population activity at each time point in the trial. Green indicates decoding from exclusively state 1 trials, and black indicates decoding from trials spanning all three states (n=7 recording sessions, mean ± s.e.m, Mann-Whitney U-test). **(E)** SVM decoding accuracy of stimulus information from PPC population activity (n=8 recording sessions, mean ± s.e.m, Mann-Whitney U-test). **(F)** SVM decoding accuracy of choice information from AC population activity (n=7 recording sessions, mean ± s.e.m, Mann-Whitney U-test). **(G)** SVM decoding accuracy of choice information from PPC population activity (n=8 recording sessions, mean ± s.e.m, Mann-Whitney U-test).

Having established that PPC activity patterns differed across performance states, we next tested whether performance state affected the encoding of task-related information. Using SVM decoding, we assessed the accuracy of predicting either auditory stimulus location or the mouse’s behavioral choice from population activity, comparing trials from the high-performance state 1 versus trials pooled across all behavioral states (See methods section “SVM decoding of state, stimulus, and choice information”). We could not train and test separate decoders for the biased states alone for two key reasons. First, mice performed the task in the high-performance state for the majority of trials (Figure 2C), and so insufficient trial numbers existed to separately train the model in the biased states for a given behavioral session. Second, by definition, the joint distribution of stimulus and choice conditions varied systematically across performance states and so it was necessary to combine the three states to fully decorrelate stimulus and choice conditions for training and testing the model. The analysis revealed that performance state had minimal impact on the quality of neural encoding in both brain regions. For stimulus decoding in AC, accuracy was similar in state 1 trials (N=7 datasets, mean = 73.4±3.05%) and all-state trials (N=7 datasets, mean=71.2±3.20%, p=0.519, Wilcoxon signed rank test, Figure 3D), indicating that AC populations maintained consistent sensory representations regardless of behavioral state. PPC stimulus decoding accuracy was also similar between trials during state 1 (N=8 sessions, mean=69.4±2.38%) and all states (N=8 datasets, mean=63.4±3.20%, p=0.246, Wilcoxon signed rank test, Figure 3E). Choice decoding from AC population activity had similar accuracy for state 1 (N=7 datasets, mean = 61.8±2.10%) versus all states (N=7 datasets, mean=62.4±3.14%, p=0.917; Wilcoxon signed rank test; Figure 3F). In contrast, choice decoding from PPC population activity was significantly higher in state 1 (N=8 sessions, mean=82.1±1.67%) compared to all states (N=8 datasets, mean=74.2±2.60%, p=0.002; Wilcoxon signed rank test; Figure 3G).

To summarize, the state decoding results demonstrate that PPC, but not AC, exhibited state-dependent population dynamics. This state-dependent information in PPC also corresponded to enhanced choice encoding during high-performance states. In contrast, AC maintained equivalent decoding performance for both stimulus and choice information regardless of performance state. Together, these results suggest that PPC, but not AC, population activity distinguishes behavioral performance state in the task.

### Single neuron encoding was state-dependent in PPC, but not AC

Neural activity in both AC and PPC is heterogeneous, affected by stimulus, choice, and various other task-relevant variables such as reward delivery, trial epoch, and the mouse’s running behavior^17,24,35,36^. To understand the relationship between these variables and neural activity in AC and PPC, we followed an encoding model approach using generalized linear models (GLMs^24^). The GLMs, which used all measured task-relevant variables as predictors of each AC or PPC neuron’s activity (See methods section “GLM encoding models” for model details), were trained and tested using trials that occurred during different behavioral performance states, as defined by the GLM-HMM above (Figure 4A). The first set of models included only trials from the high-performance state (‘state 1 encoding model’, Supplementary figure 8A), and the second set of models were trained and tested using trials from all three performance states (‘all-state encoding model’, Supplementary figure 8A). As for the stimulus and choice SVM decoding above, it was necessary to sample trials from all states to decorrelate stimulus and choice conditions for training and testing the model. We could then compare the model’s accuracy in predicting neural activity using behavioral and task variables while mice were in the high-performance state, to a mixture of performance states.

**Figure 4.**
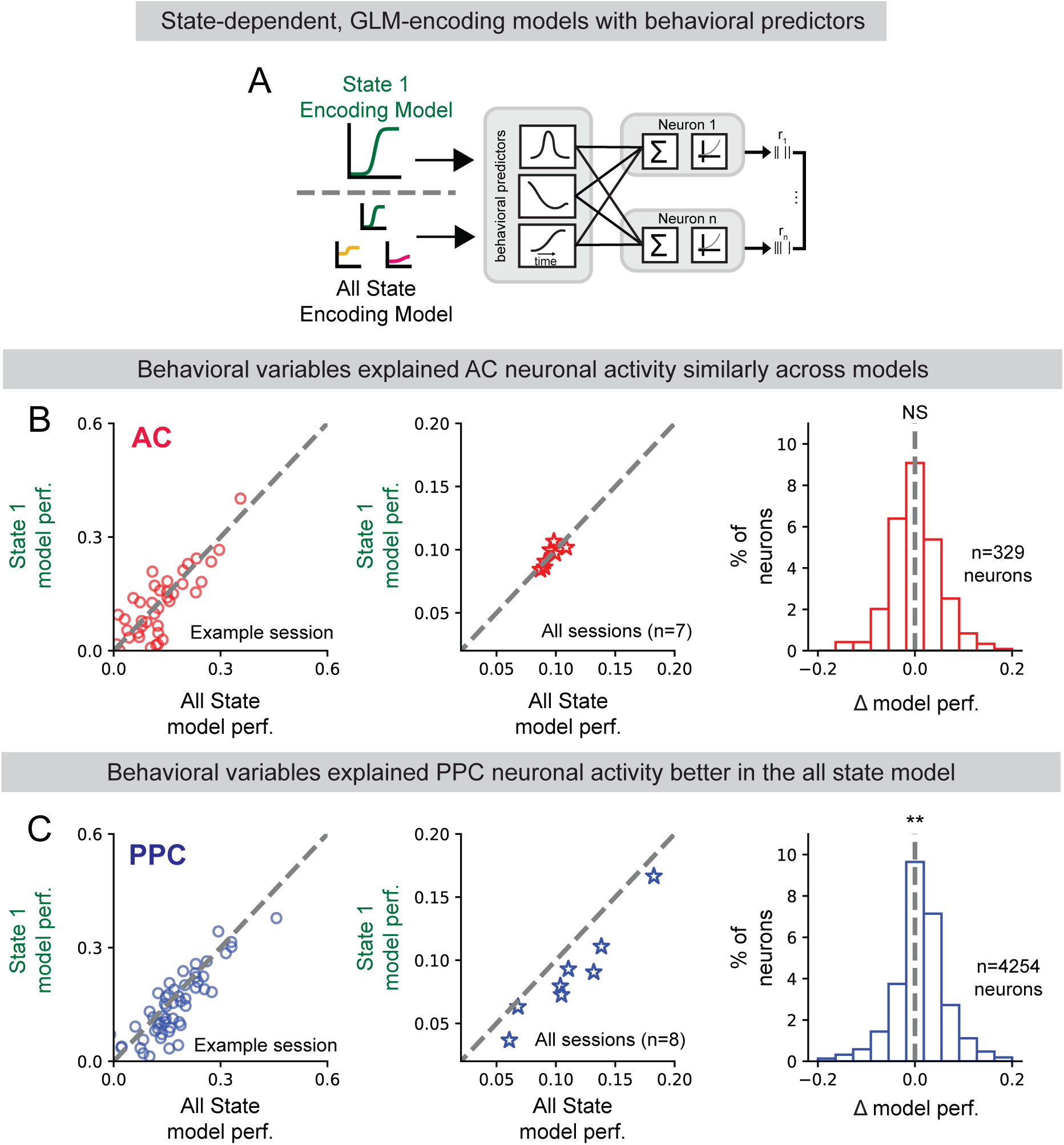
Single neuron encoding was state-dependent in PPC, but not AC. **(A)** Model schematic for two GLM-based encoding models. The state 1 encoding model was trained to predict neuronal activity given task information from exclusively high-performance state trials. The all-state model was trained to predict neuronal activity given task information from trials spanning all three states. **(B)** Prediction performance of the state 1 model and all-state model for an example session, all sessions and all AC neurons. (7 datasets, n=329 AC neurons, state 1 model mean=0.093 ± 0.008, all state model mean=0.091 ± 0.003, p=0.18, Mann-Whitney U-test). **(C)** Prediction performance of the state 1 model and all state model for an example session, all sessions, and all PPC neurons. (8 datasets, n=4254 PPC neurons, state 1 model mean=0.089 ± 0.021, all state model mean=0.112 ± 0.011, p=0.0039 Mann-Whitney U-test).

We compared the state 1 encoding model’s prediction performance (measured as the fraction of explained deviance in held-out trials) to the all-state encoding model’s prediction performance. We could then estimate the impact of biased performance states on the encoding of behavior and task-related variables in neural activity. In AC, the prediction performance of the two encoding models was similar (N=7 datasets, n=329 AC neurons; state 1 model mean=0.093±0.008; all-state model mean=0.091±0.003; p=.18, Mann-Whitney U-test, Figure 4B). Surprisingly, the activity of PPC neurons was better predicted by the all-state encoding model (N=8 datasets, n=4254 PPC neurons; state 1 model mean=0.089±0.02; all-state model mean=0.112±0.011; p=0.0039, Mann-Whitney U-test, Figure 4C). This result suggested that the activity of individual PPC neurons was more strongly related to external task variables during the biased performance states, while during optimal performance, PPC activity was more poorly explained by these measured behavioral and stimulus variables, potentially indicating that high-performance states involve greater influence from unmeasured variables.

Next, we examined the contributions of different categories of predictors in explaining neurons’ activity across states in the encoding model. To assess total weighting of each category of predictor, we summed the coefficients for each type of predictor (e.g. sound stimuli) from the model fits for each AC and PPC neuron. Sound, position/choice, and running velocity predictors were weighted similarly by both sets of encoding models for AC neurons (Sound predictors: state 1 model mean=30.5±8.03, all-state model mean=35.5±4.54, p=0.469; Position/choice predictors: state 1 model mean=9.43±1.29, all-state model mean=9.65±2.18, p=0.001; Running velocity predictors: state 1 model mean=6.70±1.09, all-state model mean=8.32±1.81, p=0.469, Mann-Whitney U Test; Supplemental figure 9A-C, top). In contrast, in PPC neurons, sound stimulus predictor coefficients were more strongly weighted in the all-state encoding model than the state-1 encoding model (state 1 model mean=15.16±0.64, all-state model mean=22.31±1.36, p=0.016, Mann-Whitney U Test, Supplemental figure 9A, bottom). Position/choice predictors’ weights were similar across states, while and running velocity predictors were more strongly weighted in the all-state models (Position/choice predictors: state 1 model mean=11.39±0.71, all-state model mean=12.93±0.71, p=0.195; Running velocity predictors: state 1 model mean=9.97±0.56, all-state model mean=12.93±1.13, p=0.008, Mann-Whitney U Test; Supplemental figure 9B-C, bottom). To summarize, the activity of AC neurons was similarly explained by stimulus and behavioral variables regardless of performance state, while the activity of PPC neurons was better explained by these variables when mice were in the biased behavioral performance states.

### Functional coupling rescued PPC encoding model performance in the high-performance state

We were initially surprised to find that task and behavioral variables more poorly predicted PPC activity when mice more accurately performed the task (Figure 4C). Previous work has shown that functional coupling, or shared variability among neurons in local populations, is stronger in association than sensory cortex especially during higher task performance accuracy and affects information coding ^24,37^. One hypothesis for why the activity of PPC neurons was more poorly explained by behavioral task variables when mice occupied the high-performance state is that functional coupling is a stronger contributor to local activity. The PPC encoding model was therefore missing a key predictor of neural activity in the high-performance state.

We first asked whether functional interactions among neurons in AC and PPC varied across performance states, by computing pairwise noise correlations, the Pearson correlation between two neurons’ trial-to-trial response variability (Supplemental figure 10). Noise correlations during the sound onset period and choice epoch in the AC population were similar between performance states (Stimulus: state 1 = 0.041±0.004, all-states = 0.039±0.005, p=0.688; Choice: state 1 = 0.029±0.002, all-states = 0.027±0.003, p=0.469; Mann-Whitney U Test, Supplemental figure 10A-B). However, noise correlations were higher during the high-performance state in PPC during the choice epoch (Stimulus: state 1 = 0.032±0.003, all-states = 0.028±0.002, p=0.383; Choice: state 1 = 0.048±0.005, all-states = 0.032±0.003, p=0.016; Mann-Whitney U Test, Supplemental figure 10C-D). This indicated that shared variability across PPC neurons was more pronounced during periods of high task performance and could therefore be the missing predictor of neuronal activity in the encoding model explaining activity in PPC in the high-performance state (Figure 4).

To test this hypothesis and measure functional coupling in PPC population activity across performance states, we modified our encoding models to predict a given neuron’s activity based on task information and the activity of the other neurons in the population (Figure 5A). We extracted the first five principal components (PCs) of single trial population activity concatenated across all trials (excluding the activity of the predicted neuron) and convolved these PCs with gaussian basis functions to model correlations across time ^24^. We refit both the all-state and state-1 encoding models for each AC or PPC neuron with these ten additional ‘functional coupling’ predictors and compared prediction performance with and without population activity to measure the contribution of functional coupling to each neuron’s activity that cannot be explained by behavioral and task-related variables alone. For AC neuronal activity, adding functional coupling predictors had no significant effect on prediction performance for both the state 1 and all-state models (N=7 datasets, n=329 neurons, State 1 model: behavior only model mean = 0.093±0.009, behavior + coupling model mean = 0.095±0.011, p=.812; All-state model: behavior only model mean = 0.091±0.004, behavior + coupling model mean = 0.087±0.006, p=.469; Mann-Whitney U-test, Figure 5B-C). However, adding functional coupling predictors to the state 1 encoding model for PPC neurons significantly increased prediction performance in comparison to the state 1 model without population activity (N=8 datasets, n=4254 neurons, State 1 model: behavior only model mean = 0.089±0.007, behavior + coupling model mean = 0.122±0.004, p=0.039, Mann-Whitney U-test, Figure 5D). For the all-state encoding model, adding functional coupling predictors had no significant effect on prediction performance of PPC neuronal activity (N=8 datasets, n=4254 neurons, State 1 model: behavior only model mean = 0.112±0.007, behavior + coupling model mean = 0.118±0.003, p=0.547, Mann-Whitney U-test, Figure 5E). Additionally, an examination of the model weights for the functional coupling predictors for each model revealed that coupling weights were larger in the state-1 model than in the all-state model, only for PPC neurons (AC: State 1 model mean = 6.97±0.6, All state model mean = 6.67±0.9, p=.938; PPC: State 1 model mean = 16.83±1.19, All state mean = 8.80±1.13, p=0.008, Mann-Whitney U-test, Supplemental figure 11D).

**Figure 5.**
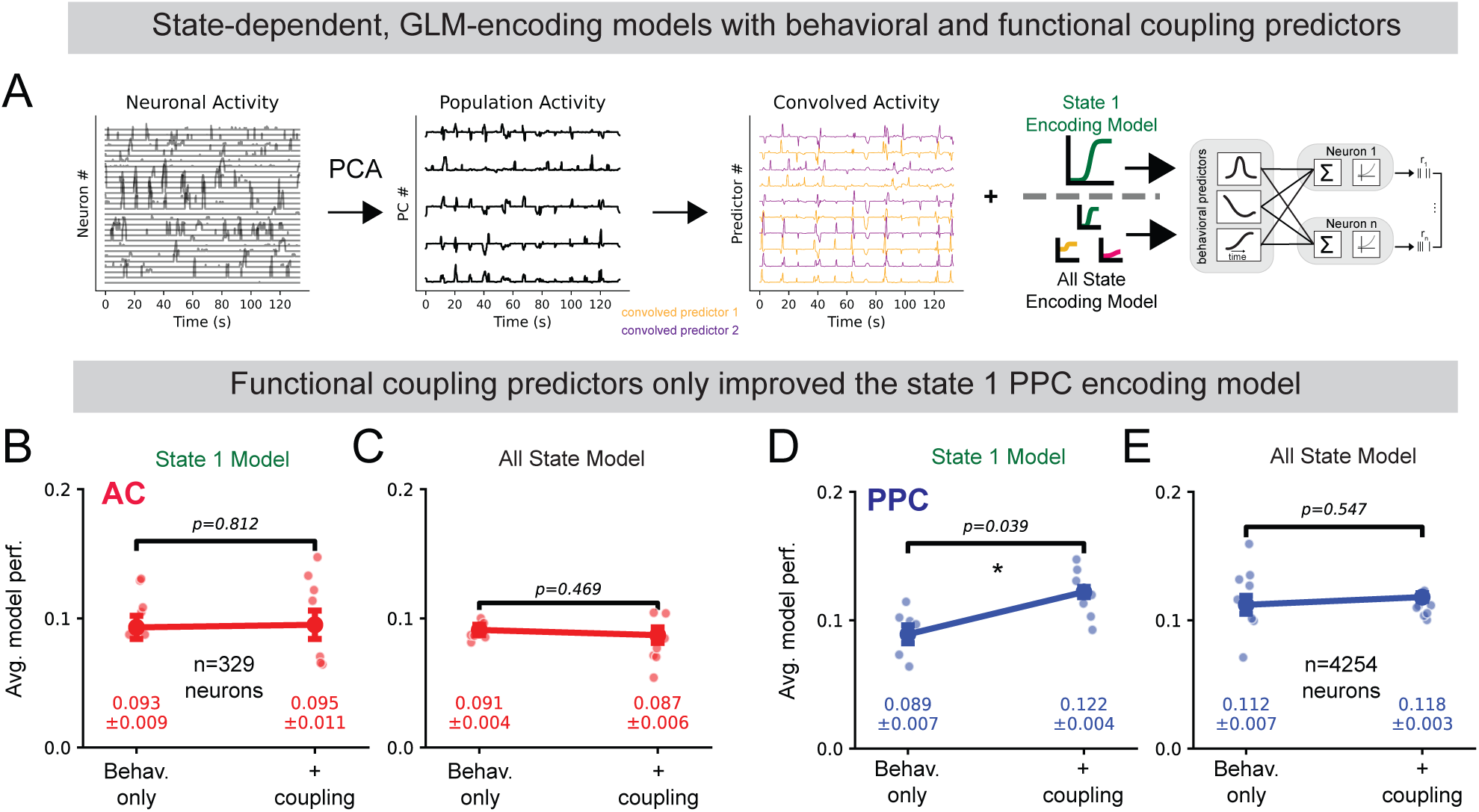
Functional coupling rescued PPC encoding model performance in the high-performance state. **(A)** Schematic of the PCA based population activity functional coupling predictors used in the encoding models. Single trial neuronal activity is concatenated across all trials in the training set and is used to compute PCA. **(B)** Comparison of average state-1 model performance with and without functional coupling predictors for AC neuronal activity (7 datasets, n=329 neurons, Behavior only model mean = 0.093±0.009, Behavior + coupling model mean = 0.095±0.011, p=.812, Mann-Whitney U-test). **(C)** Comparison of average all-state model performance with and without functional coupling predictors for AC neuronal activity (7 datasets, n=329 neurons, Behavior only model mean = 0.091±0.004, Behavior + coupling model mean = 0.087±0.006, p=.469, Mann-Whitney U-test). **(D)** Comparison of average state-1 model performance with and without functional coupling predictors for PPC neuronal activity (8 datasets, n=4254 neurons, Behavior only model mean = 0.089±0.007, Behavior + coupling model mean = 0.122±0.004, p=.039, Mann-Whitney U-test). **(E)** Comparison of average all-state model performance with and without functional coupling predictors for PPC neuronal activity (8 datasets, n=4254 neurons, Behavior only model mean = 0.112±0.007, Behavior + coupling model mean = 0.118±0.003, p=.547, Mann-Whitney U-test).

## Discussion

Animals flexibly adjust their decision-making strategies to optimize performance in dynamic environments, yet the neural processing implications of these behavioral state transitions remain poorly understood. Here, we combined computational modeling of choice behavior in an auditory decision-making task with previously published neural population recording data^24^ and newly collected causal manipulations. Our goal was to investigate how cortical circuits are impacted by changes in behavioral performance state during the task. Similar to the performance states identified during visual decision-making ^4–6^, we identified one high performance state and two biased states in which mice were biased in their choice behavior toward either left or right choices (Figure 2B, C). We related population activity within the posterior parietal cortex (PPC) and the primary auditory cortex (AC) to these behavioral performance states, and found that behavioral performance state was linked to different population activity states in PPC, but not in AC. Crucially, inactivation of PPC shifted mice into the biased performance state, establishing a causal link between PPC activity and behavioral performance state in the task.

Previous studies assessing the causal requirement of PPC in perceptual decision-making tasks have been contradictory^27,28,32,38,39^, yet we found that PPC activity was required for accurate performance of the auditory localization task in VR (Figure 1). The causal requirement of PPC for decision-making likely depends on the task paradigm. For example, rats making auditory decisions did not require PPC activity for accurate task performance^18^, nor did mice making decisions based on well-learned stimulus sets^29^. In these paradigms, sensory cues were presented and animals responded to the cues in separate task epochs. In our task, mice must localize a sound while navigating down a virtual hallway and then report the decision by turning at the T-maze intersection. This task design naturally increases the cognitive load on the mouse, while previous studies used paradigms with stationary frameworks for reporting choices, such as headfixed mice responding by licking^29^. Additionally, in our experiments we found that the causal requirement of PPC persisted for multiple sessions, as performance was degraded on every session in which we inactivated PPC (Supplemental Figure 4). This result reinforces the requirement of PPC for continued timescales in our paradigm.

In addition to its causal requirement for mice performing the task, population activity patterns in PPC strongly distinguished mouse’s behavioral performance state in two key ways (Figures 3-5). First, even when controlling for the mouse’s left-right choices (a strong driver of PPC activity) we were able to predict whether the mouse was performing the task in a particular trial in the high performance or in one of the biased states using PPC, but not AC population activity patterns (Figure 3C). Second, an encoding model’s ability to accurately predict single neuron activity in PPC hinged on including the activity of each neuron’s local population as predictors (‘functional coupling’), but only in the high-performance state (Figure 5D-E), suggesting that population activity is more coordinated in the high-performance state. These state-dependent population activity patterns in PPC were also associated with enhanced choice decoding during high-performance periods (Figure 3G), while stimulus encoding remained stable across states (Figure 3E). This pattern suggests a functional advantage for state-dependent activity patterns in PPC, rather than a simple dissociation between state representation and information encoding.

The circuit mechanisms underlying state-dependent changes in ‘functional coupling’ are likely to include a combination of brain wide changes in neuromodulation, and changes in specific longe-range or local circuit interactions. For example, the activity in neuromodulatory nuclei such as the locus coeruleus and nucleus basalis is correlated with task engagement, motivation, and arousal ^1,40–42^. Changes in neuromodulatory tone within the PPC or in its inputs from other brain areas such as the prefrontal cortex or subcortical regions could then alter the activity levels of specific neuron types, such as inhibitory subtypes^43–45^ to modulate local population activity dynamics. Increased efficacy of PFC-PPC recurrent interactions^46–48^ could indirectly drive increases in functional coupling within PPC, as could increased efficacy of local excitatory recurrent synapses within PPC. Higher functional coupling can provide a computational advantage by creating more consistent and reliable population signals that facilitate accurate decoding by downstream brain areas^24,37^. Future studies employing causal manipulations of these potential sources will be required to reveal the underlying mechanisms contributing to the level of local functional coupling.

Unlike PPC, activity in auditory cortex (AC) was unaffected by performance state in this task. One hypothesis is that the shifts from high to biased performance state could be related to shifts in the arousal state of the animal. A rich literature has established relationships between arousal state and stimulus coding, firing rates, and shared variability in auditory cortex^49–52^ and so the current results may seem contradictory. A few possibilities could explain this inconsistency. We did not monitor pupil diameter, so we cannot determine whether arousal states shifted systematically between states. However, the psychometric curves measured during the biased performance state trials suggest that although mice were biased in reporting choices, they remained engaged in performing the task. Additionally, our task design requires mice to be sufficiently aroused to voluntarily progress through the T-maze by running on the spherical treadmill, suggesting that our mice were consistently in a relatively aroused state where small differences in arousal across GLM-HMM states may not be wide enough to differentially drive AC activity patterns. Running behavior is often used as a proxy for arousal level in mice^50,51,53^, and mice ran through T mazes in both high and biased performance states in our task, with even higher speeds in the biased states (Supplementary Figure 6B).

More fundamentally, the differential impact of behavioral performance state on PPC versus AC likely reflects their distinct functional roles in decision-making. While AC did encode task information beyond sensory stimuli (Supplementary Figure 9), we found that AC maintained relatively stable stimulus representations regardless of behavioral strategy (Figure 3). In contrast, PPC integrates multimodal sensory evidence with internal goals, expectations, and cognitive strategies^18,23,30,32,54–56^. This positions PPC to be more sensitive to the types of strategic shifts captured by our GLM-HMM states. Future studies could test this hypothesis by examining whether manipulations that specifically alter cognitive control, rather than arousal or sensory processing, preferentially impact PPC versus AC activity patterns during decision-making tasks^43–46^.

Notably, task and movement-related variables better explained PPC activity in biased performance states compared to high-performance states, aligning with evidence that uninstructed movements and spontaneous behavioral fluctuations are inversely correlated with cognitive performance^54,55^. Recent studies across multiple sensory and decision-making tasks have demonstrated that periods of poor performance are characterized by increased influence of task-irrelevant movements on cortical activity^57–59^. Our results suggest that during biased performance states, PPC activity becomes more correlated with these behavioral variables, potentially reflecting a loss of top-down cognitive control. These findings also align well with emerging evidence that spontaneous, task-independent activity dynamics across cortical areas influence behavioral performance independent of sensory processing^7,60^. Our results extend this framework by demonstrating that PPC not only exhibits state-dependent population dynamics, but that these dynamics are related to specific behavioral strategies during decision-making.

### Limitations of the study

A limitation to our approach of comparing cortical encoding across performance states is that mouse behavior in biased states is, by definition, biased toward one choice direction in a particular state. Training and testing models to compare encoding and decoding of variables such as choice across behavioral states then required that we subsample trials to ensure that variables such as stimulus, choice, and behavioral state are separable. Additionally, we measured various behavioral features during task performance such as running velocity, sensory stimulus identity and timing, behavioral choice, and reward delivery. We used these variables in our GLM-HMM modeling and GLM encoding analyses, and inferred latent states related to behavior, which we connected to neural activity patterns. However, we did not measure arousal metrics such as pupil size.

An additional limitation concerns the interpretation of choice-related neural activity in PPC. Because behavioral choice and running kinematics are inherently linked in our T-maze paradigm, we cannot fully disentangle neural activity related to motor execution from activity more directly linked to the decision-making process itself. While this link allows us to decode the animal’s evolving choice commitment as they integrate sensory information, it also means that the enhanced choice decoding accuracy observed in state 1 could partially reflect differences in movement kinematics rather than purely cognitive decision-related signals.

Finally, several limitations are inherent in the calcium imaging approach. Because of their different optical angles, calcium imaging data from AC and PPC could not be collected simultaneously. The slow timescale of calcium imaging compared to electrophysiology experiments limits our measurement of functional coupling to longer timescales (10s to hundreds of milliseconds). We take this into account when comparing across both regions and recognize that correlations between the two populations could also be modulated by transitions in performance state.

## Acknowledgements

We thank Chengcheng Huang, Jennifer H. Fang, and members of the Runyan lab for comments on the manuscript. We thank Christopher Harvey (Harvard Medical School) in whose laboratory a portion of the neural data was collected. We thank the GENIE project (Janelia) for making GCaMP sensors available for use. This work was supported by the NIH Predoctoral Training Grant in Basic Neuroscience T32NS126122-02, Pew Biomedical Scholars Program, the Searle Scholars Program, the Klingenstein-Simons Fellowship Award in Neuroscience, and NIH grants NIMH DP2MH122404 and NINDS R01NS121913.

## Author Contributions

A.C.B. and C.A.R. designed the experiments, A.C.B., J.P.M., R.M.C., M.J.L., J.N.L. and C.A.R. performed the experiments, A.C.B. performed analyses, and A.C.B. and C.A.R. wrote the paper with input from all authors.

## Data and code availability

Analysis code is deposited and publicly available at: https://github.com/acbandi213/Bandi---Runyan-2024.git

## Experimental model details

Imaging data were previously collected from five male C57BL/6J mice^24^ (The Jackson Laboratory), and new imaging data was collected from two female transgenic mice that were bred to express GcAMP6s in cortical excitatory neurons (B6.Cg-Tg(Camk2a-tTA)1Mmay/DboJ x B6;DBA-Tg(tetO-GCaMP6s)2Niell/J) that were ∼ 7 weeks old at the initiation of behavior task training. Optogenetic inactivation and behavioral data was collected from eight C57BL/6J mice. 5 male mice were injected with stGtACR2 and used for optogenetic inactivation, and 3 without stGtACR2 injection were used for control inactivation (2 male and 1 female).

## Method details

In this study, we performed novel behavioral, optogenetic inactivation and calcium imaging experiments. We also performed an independent analysis of publicly available mouse behavioral and calcium imaging experiments described previously^24^. A summary of the experimental procedures is provided here, and full detailed procedures related to neuronal recordings can be found in the previously published work. All experimental procedures were approved by the University of Pittsburgh Institutional Animal Care and Use Committee and Harvard Medical School Institutional Animal Care and Use Committee.

### Sound localization task

Head-restrained mice ran on a spherical treadmill to control movement through a virtual reality T-maze, which was projected on a screen in front of the mouse. The virtual T-maze was constructed using the Virtual Reality Mouse Engine (ViRMEn^61^) in MATLAB (v2011a). As mice ran down the stem of the T-maze, sound stimuli were 1-2 s dynamic ripples, delivered from one of eight possible locations (−90°, −60°, −30°, −15°, +15°, +30°, +60° and +90°; from the previously collected datasets^24^), or from two possible locations (−90° and +90°; used for newly collected datasets in transgenic animals used for imaging experiments and for animals used in optogenetic experiments), using speakers centered around the mouse’s head. The stimuli began 10cm into the maze and repeated after a 100-ms gap until the mouse reached the T-maze intersection. Mice were required to report the left-right category of the stimulus location by turning in that direction into either the left or right arm at the T-maze intersection. Correct decisions resulted in delivery of a 4 μl sugar water and a ‘reward tone’, while incorrect decisions resulted in a ‘no-reward tone’. Following a correct choice, there was a 3 s inter-trial interval (ITI), and following an incorrect choice there was a 5 s ITI.

### Optogenetic inactivation of parietal cortex

For optogenetic inactivation experiments, mice performed a simplified version of the sound localization task where only −90° and +90° speaker locations were used as stimuli. This simplified task was used to ensure enough inactivation trials per condition. We bilaterally injected a soma-targeted anion channelrhodopsin (stGtACR2) at a depth of 200 µm in PPC to target excitatory neurons via the CamKIIa promotor. Three mice were implanted with fiber optic canula above the virus injection which were used to deliver 470nm light at 1.5mW power. Two mice were implanted with cranial windows above the virus injection and the 470nm light was placed directly on the cranial window. This approach allows us to inactivate PPC excitatory activity, by activating neurons expressing stGtACR2 using 470nm light at specific time points in the task on a subset of trials. We bilaterally inactivated the PPC from stimulus onset to the stem of the T-maze prior to the mouse turning into the arm to report its choice. Inactivation occurred on 25% of trials of a specific context. For control inactivation experiments, mice (N=3) without stGtACR2 injection were implanted with fiber optic canula and 470nm light at 1.5mW power was delivered.

### In vivo calcium imaging

Imaging was performed on alternating days from the AC and PPC on the left hemisphere of the animal (PPC centered at 2 mm posterior and 1.75 mm lateral to bregma; AC centered at 3.0 mm posterior and 4.3 mm lateral to bregma). In each session, ∼50 neurons (range, 37–69) were simultaneously imaged using a two-photon microscope (Sutter MOM) operating at a 15.6-Hz frame rate and at a resolution of 256 × 64 pixels (∼250 μm × 100 μm). ScanImage (version 3, Vidrio Technologies) was used to control the microscope. Imaging data were acquired at depths of between 150 and 300 μm, corresponding to layers 2/3. Seven AC and eight PPC fields of view from five mice were analyzed.

### Quantification and statistical analysis

#### GLM-HMM modeling of behavioral performance

To quantify transitions between discrete decision-making states within a single behavioral session, we used a hidden Markov model with Bernoulli Generalized Linear Model observations (GLM-HMM) based on a modified version of the SSM python package. The model is defined by a transition matrix containing a fixed set of transition probabilities: z ∈{1,…,K}, and a vector of GLM weights for each state. Each GLM has a unique set of weights w*_k_* that maps external covariates to the probability of choice for each of the *k* states. We coded the external covariates on each trial as follows: (1) the signed location of the auditory stimulus on the present trial; (2) the binary choice (1 for right or 0 for left) made by the mouse on the previous trial; (3) whether the previous trial was correct or incorrect (1 or 0); (4) and a constant offset or bias. The output for each trial was a value of 1 or 0 depending on whether the mouse turned right or left.

We first fit the GLM-HMM to behavioral data from mice performing the eight-location behavioral task^24^ (N=5 mice, 17602 trials; Figure 1A-C, Figure 2A-E), using the expectation-maximization (EM) algorithm, again using the SSM python package. The GLM-HMM state was inferred using the posterior probabilities calculated from the preceding trials and the state transition matrix. To select the number of latent states in the model (K) we performed cross-validation of the behavioral data, which revealed that three states allowed the model to plateau in likelihood, calculated via maximum likelihood estimation (MLE). We also measured choice prediction accuracy for the three state GLM-HMM by using the weights of the inferred state to predict the choice on that trial, which is compared to the empirical data thus determining model prediction accuracy. Following this optimization of states, we then fit a single three state GLM-HMM to the observations and inputs concatenating all sessions from a single subject and again inferred the state occupation estimates using the posterior probabilities calculated from the preceding trials and the state transition matrix. We found that the three states were consistent across all the subjects used in this study and each three state GLM-HMM fit to each individual animal had high log-likelihood and predictive accuracy on held-out test data. For all further analyses, we set an 80% state probability criterion for inclusion of a trial with a performance state, and discounted trials that did not meet the criterion. This same approach was used to fit the GLM-HMM to behavioral data in which PPC was optogenetically inactivated (N=5 mice, 5351 trials) and we found that the same three states were consistent across all the subjects.

#### SVM decoding of state, stimulus, and choice information

For population decoding of performance state information, we used a SVM decoder with a linear kernel (C=100, gamma=0.1, identified via best estimate grid search) based on the sklearn.svm python package. Equal numbers of state 1 trials and non-state 1 trials (combining state 2 and 3) were selected and then balanced to have equal representation of trials from each stimulus and choice combination (left stimulus left choice, right stimulus right choice, left stimulus right choice, right stimulus left choice), ensuring that any state differences in neural activity could not be attributed to differences in sensory input or motor output. We trained the SVM decoder to classify a trial as state-1 or not state-1 (combining states 2 and 3) from AC or PPC neuronal population activity during that trial and calculated the decoding accuracy using five-fold cross validation. We pooled population activity from distinct time points in the trial. The first time point is aligned to the onset of the auditory stimulus and goes 3 seconds post stimulus onset. The second time point is aligned to turn onset and uses 3 seconds pre turn onset. We concatenated these two temporal bins. State, stimulus and choice information was then classified using an SVM decoder across these pooled windows.

For comparing the population decoding of stimulus and choice information across performance states, we first separated trials based on the state as identified by the three state GLM-HMM. Due to the higher percentage of state-1 trials in comparison to state 2 and 3 trials, particularly in the behavioral sessions in which imaging was performed, we separated data into two groups for this decoding approach. The first group (‘Exclusively State 1’) included data exclusively from state-1 trials and the second group (‘All States’) included data from state 2 and state 3 trials supplemented with subsampled state-1 trials to ensure a similar amount of trials for comparisons between groups. In an area like PPC where neurons strongly encode choice direction, direct comparison of the three states could lead to trivial conclusions based on the differential choice outcomes in the left and right biased states. It was therefore crucial to balance all possible combinations of stimulus and choice directions to decouple these variables and balance their weighting in the training datasets. We accomplished this by combining the two biased performance states, balancing their trial numbers, in the training and testing datasets for our models. Furthermore, because the mice were trained to become experts at the auditory decision-making task, they tended to predominantly perform the task in the optimal state. The resulting imbalance, in the number of trials performed across the three states, left too few trials to directly compare population coding between biased and high-performance states, as these analyses require large numbers of trials to train encoding models and test their prediction performance. We therefore compared population coding in high performance (state 1) vs ‘all’ states, combining the two biased states and supplementing with trials during high performance. For each group, within the training dataset (70% of trials), we ensured that cross-validation folds had equal number of trials from each stimulus and choice combination (left stimulus left choice, right stimulus right choice, left stimulus right choice, right stimulus left choice). The test dataset (30% of trials) also had a similar distribution of trial conditions and was left out of the fitting procedure. All decoders were built and trained using the sklearn.svm python package. Independent SVMs were trained and tested at each time point, and decoding accuracy was expressed as the proportion of correct classifications across the folds of cross-validation.

#### Balancing trial conditions to decorrelate stimulus, choice, and behavioral performance state

To ensure that differences in neural activity between behavioral states could not be attributed to confounding factors such as stimulus or choice responses, we implemented a rigorous trial balancing scheme. Equal numbers of state 1 trials and non-state 1 trials (combining state 2 and 3 trials) were selected from the initial trial distribution. Given the imbalanced nature of the original dataset, this selection required downsampling from the more abundant state 1 trials to match the combined count of state 2 and 3 trials. Within each behavioral state category, trials were then systematically balanced to achieve equal representation across all four possible stimulus-choice combinations: left stimulus with left choice, right stimulus with right choice, left stimulus with right choice (error trials), and right stimulus with left choice (error trials). This balancing was applied to both the state 1 decoder training subset (∼80% correct/20% incorrect performance) and the comprehensive dataset used for the all-state SVM decoder or encoding models. The balanced design resulted in equal trial counts for each stimulus-choice combination within both state categories. This approach effectively decorrelated behavioral performance state from both sensory and choice patterns, ensuring that any observed differences in neural decoding accuracy or encoding model predictions between states could be attributed specifically to state-dependent changes in neural representations rather than systematic biases in task conditions or behavioral responses.

#### GLM encoding models

Our GLM based encoding models allowed us to assess, for each single AC and PPC neuron, the time-dependent effects of various task and behavioral variables on neuronal activity during single trials in a specific recording session. We extend the approaches taken in ^20,24,62,63^ to account for performance state by using separate encoding models for exclusively state-1 trials and another model for trials spanning all states. Again, due to the higher percentage of state-1 trials in comparison to state 2 and 3 trials, particularly in the behavioral sessions in which imaging was performed, we separated data into two groups for this encoding approach. The first group included data exclusively from state-1 trials and the second group included data from state 2 and state 3 trials supplemented with subsampled state-1 trials to ensure a similar amount of trials for comparisons between groups. For each group, within the training dataset (70% of trials), we ensured that cross-validation folds had equal number of trials from each stimulus and choice combination (left stimulus left choice, right stimulus right choice, left stimulus right choice, right stimulus left choice). The test dataset (30% of trials) also had a similar distribution of trial conditions and was left out of the fitting procedure. We used a Bernoulli GLM to weight task variables or task variables + functional coupling variables (principal components of population activity) in predicting a single neuron’s binary activity.

#### Task-related model predictors

A total of 419 task-related predictors (420 when including a constant predictor corresponding to the average activation probability of each individual neuron) were used in all encoding models. The behavioral variables included the running velocity on the pitch and roll axes of the spherical treadmill, x and y position, onset times and locations of sound stimuli, view angle of the mouse in the VR maze, turn direction, and reward and error signal delivery times. A detailed description of the selection, construction, and normalization parameters of task-related predictors using various sets of basis-functions can be found in Runyan et al., 2017.

#### Functional coupling model predictors

We developed encoding models with functional coupling predictors to compare the dependence of each neuron’s activity on task-related information correlates and the activity of the other neurons in the population. Previous work has used the relative spike rate of each other neuron excluding the neuron being fitted and convolved the spike rate with boxcar functions, however we took a dimensionality reduction approach to reduce the number of total coupling predictors. We first excluded the neuron being fit by the encoding model and performed principal component analysis (PCA) along ∼1 second time bins on the matrix of spiking activity of all other neurons in the local population for that imaging session using the sklearn.decomposition.pca python function. We then took the first five principal components (PCs), which accounted for ∼65-70% of the overall variance in the population activity. We then maximum-normalized and z-scored the PCs and convolved the PCs using two evenly spaced Gaussian basis functions extending ∼120ms second forwards and backwards in time which yielded 10 total ‘functional coupling’ predictors.

#### GLM fitting procedure

All task-related information and functional coupling predictors were maximum-normalized, and z-scored before fitting each encoding model. We fitted the GLMs to each single neuron’s activity using the GLM_tensorflow_2 python package. We used an elastic net regularized GLM which interpolates between L_1_ and L_2_ regularization penalties based on the interpolation parameter α. We used α = .95 to allow for a relatively small number of predictors to be selected by the model. Within the training dataset (70% of trials), cross-validation folds were balanced with a structured distribution of trials from each stimulus and choice combination. The test dataset (30% of trials), also containing a similar distribution of trial conditions, was left out of the fitting procedure entirely, and was used only for testing the model performance. Each model was thus fitted and tested on entirely separate data, removing over-fitting concerns. This train and test procedure was repeated ten times, with random subsamples of the data included in train and test segments.

#### GLM model performance

Model performance was quantified by computing the fraction of explained deviance of the fitted model by comparing the deviance of the fitted model with the deviance of null model of each neuron’s activity that used a single constant parameter. This null model lacked time-varying task predictors or functional coupling predictors. Thus, the fraction of explained deviance was calculated as ((null deviance – fitted model deviance) / null deviance). All deviance calculation were calculated on the test dataset for all folds of the encoding models for each neuron.

#### Pairwise noise correlations

Pairwise noise correlations were calculated based on trial-to-trial fluctuations around mean sound-evoked responses during distinct task epochs. We defined two temporal epochs for analysis: the stimulus epoch (from sound onset to 1s after onset, with activity aligned to first sound onset) and the choice epoch (from 1s before turn onset to turn onset, with activity aligned to turn onset). For each epoch and each of the eight possible sound locations, we calculated the mean activity for each neuron and binned the activity by bin size (e.g., 100ms bins). For each neuron, we subtracted the corresponding mean evoked responses from single trial activity within each epoch. For each pair of neurons, we computed the Pearson correlation coefficient between these binned, mean-subtracted activity timeseries within each epoch using the np.corrcoef python function. Noise correlations were calculated separately for trials classified as State 1 (high-performance) and for all trials combined (All States), allowing comparison of correlation structure across different behavioral performance states and task epochs.

#### Histology

After all optogenetic inactivation sessions had been acquired, each mouse was transcardially perfused with saline and then 4% paraformaldehyde. The brain was extracted, cryoprotected, embedded, frozen, and sliced. Once slide mounted, we stained brains with DAPI to be able to identify structure. We used anatomical structure to verify the locations of our stGtACR2 injections and fiber optic implants in PPC.

---

**Figure S1.**
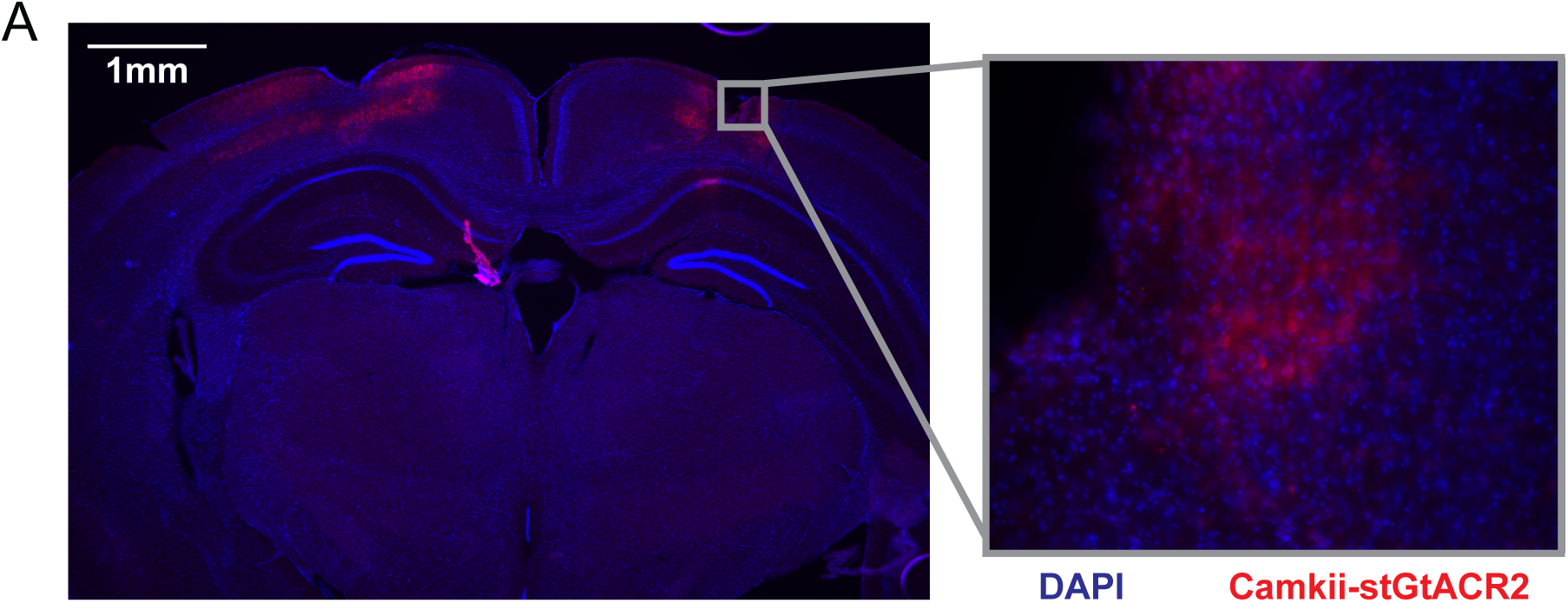
Histological confirmation of stGtACR2 expression. **(A)** Expression of AAV-CkIIa-stGtACR2 PPC neurons and fiber tracks from chronically implanted bilateral optical fiber implants (fixed tissue, DAPI in blue). Inset is zoomed 20x.

**Figure S2.**
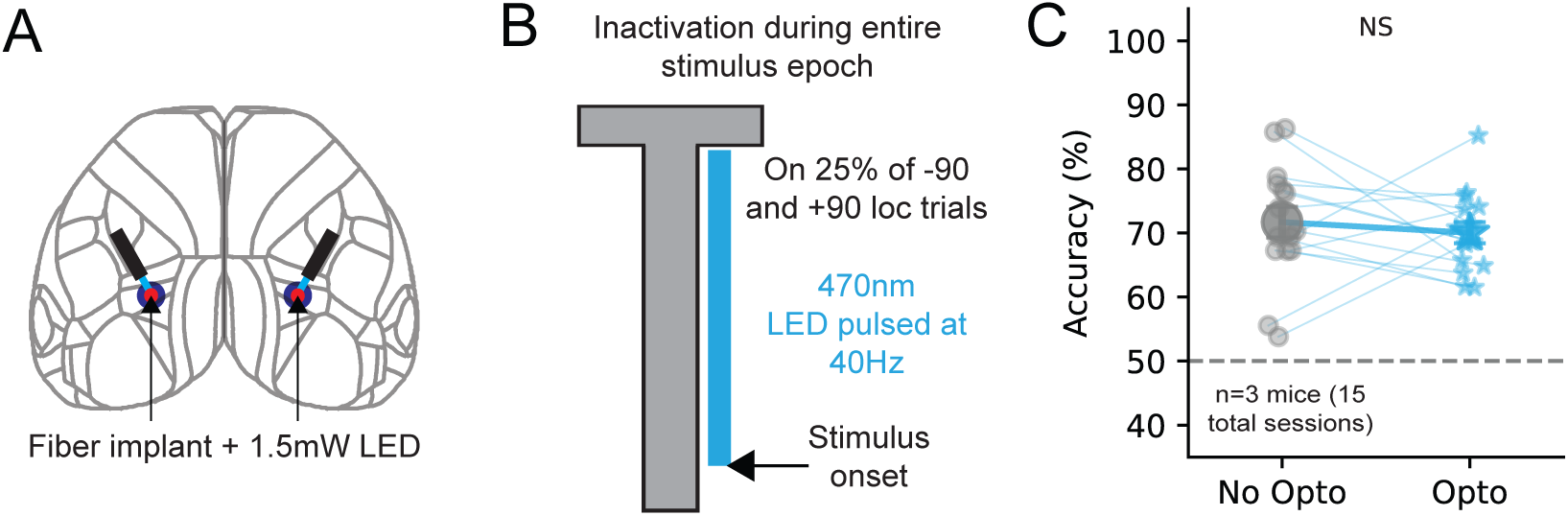
Control inactivation of PPC during the auditory decision-making task. **(A)** Schematic showing the location of optical fiber implant in PPC without injection of stGtACR2. **(B)** Schematic showing 470nm LED from stimulus onset to the stem of the t-maze on 25% of −90 and +90 sound location trials. **(C)** Effect of control LED activation of PPC on task accuracy (n=3 mice, 15 total sessions, No opto mean = 71.3 ± 4.2, Opto mean = 70.6 ± 3.1, p= 0.48, Wilcoxon signed rank test).

**Figure S3.**
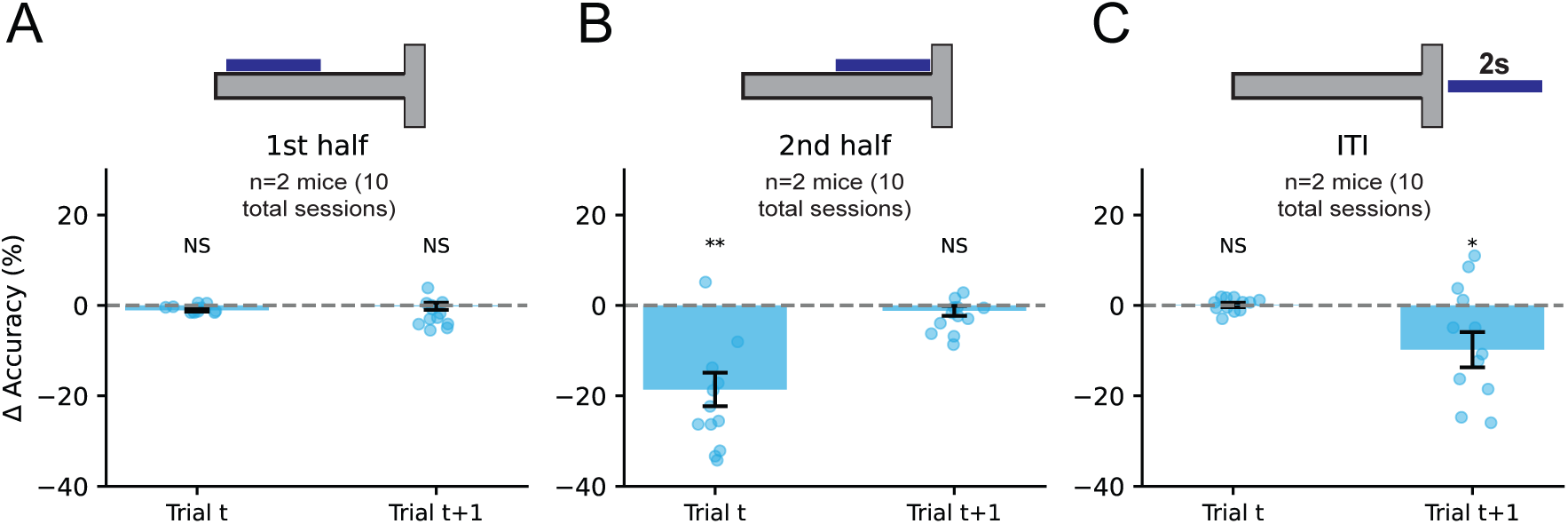
Epoch-based inactivation of PPC activity. **(A)** Change in task accuracy when PPC was inactivated during the first half of the t-maze (trial t mean = −1.1 ± 0.8, p = 0.561; trial t+1 mean = −0.2 ± 0.8, p = 0.367; Wilcoxon signed rank test). **(B)** Change in task accuracy when PPC was inactivated during the second half of the t-maze. Second half inactivation resulted in significant decrease in task accuracy during trial t. (trial t mean = −18.6 ± 3.7, p = 0.006; trial t+1 mean = −1.2 ± 1.1, p = 0.4; Wilcoxon signed rank test). **(C)** Change in task accuracy when PPC was inactivated for 2 seconds during the intertrial interval. (trial t mean = 1.1 ± 0.5, p = 0.8; trial t+1 mean = −9.8 ± 3.9, p = 0.021; Wilcoxon signed rank test).

**Figure S4.**
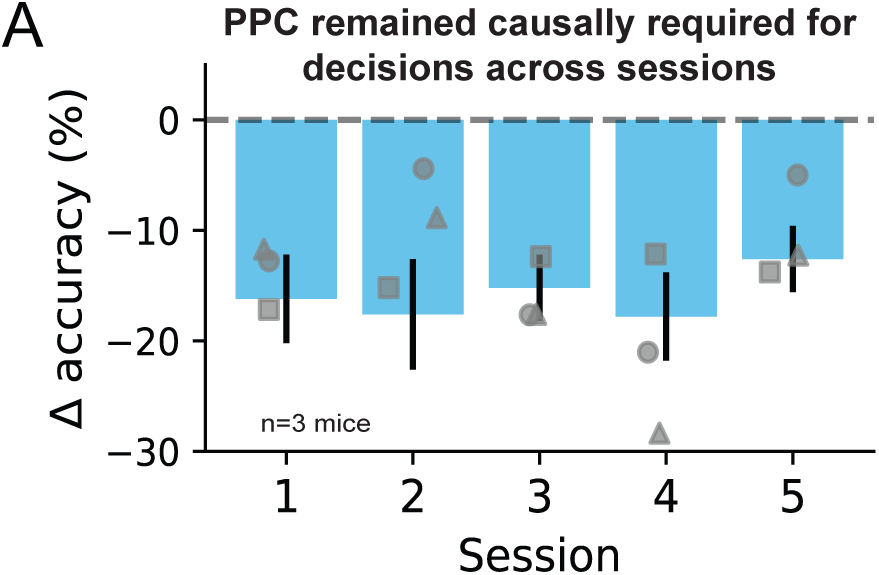
Parietal cortex remained causally required across multiple sessions. **(A)** PPC was inactivated for five repeated sessions. Decreases in task accuracy persisted across all five sessions.

**Figure S5.**
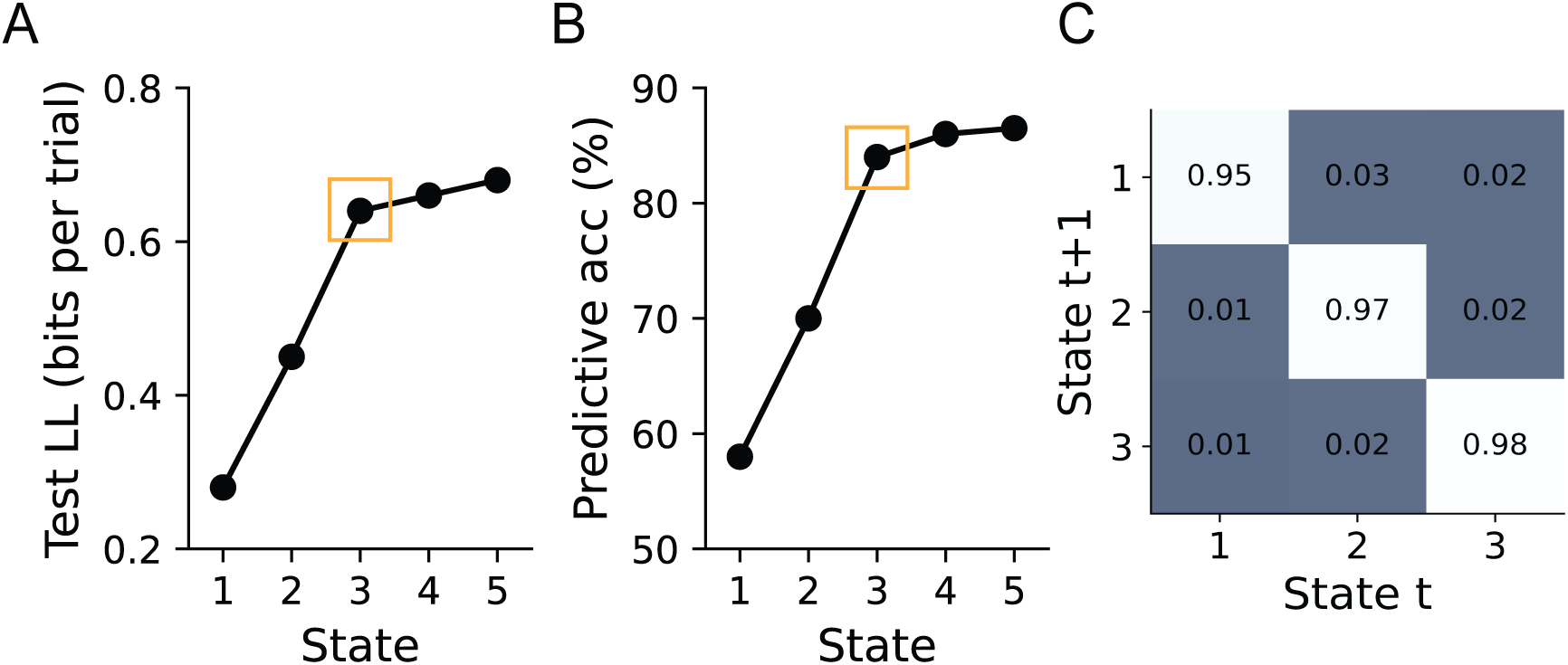
GLM-HMM cross validation. **(A)** Model comparison of GLM-HMMs with different number of latent states using test log likelihood (in bits per trial) from five-fold cross validation. **(B)** Predictive accuracy (percentage of held out test trials in which the model correctly predicted the mouse’s choice) of GLM-HMMs with different number of latent states. **(C)** Inferred transition probability matrix for global three state GLM-HMM.

**Figure S6.**
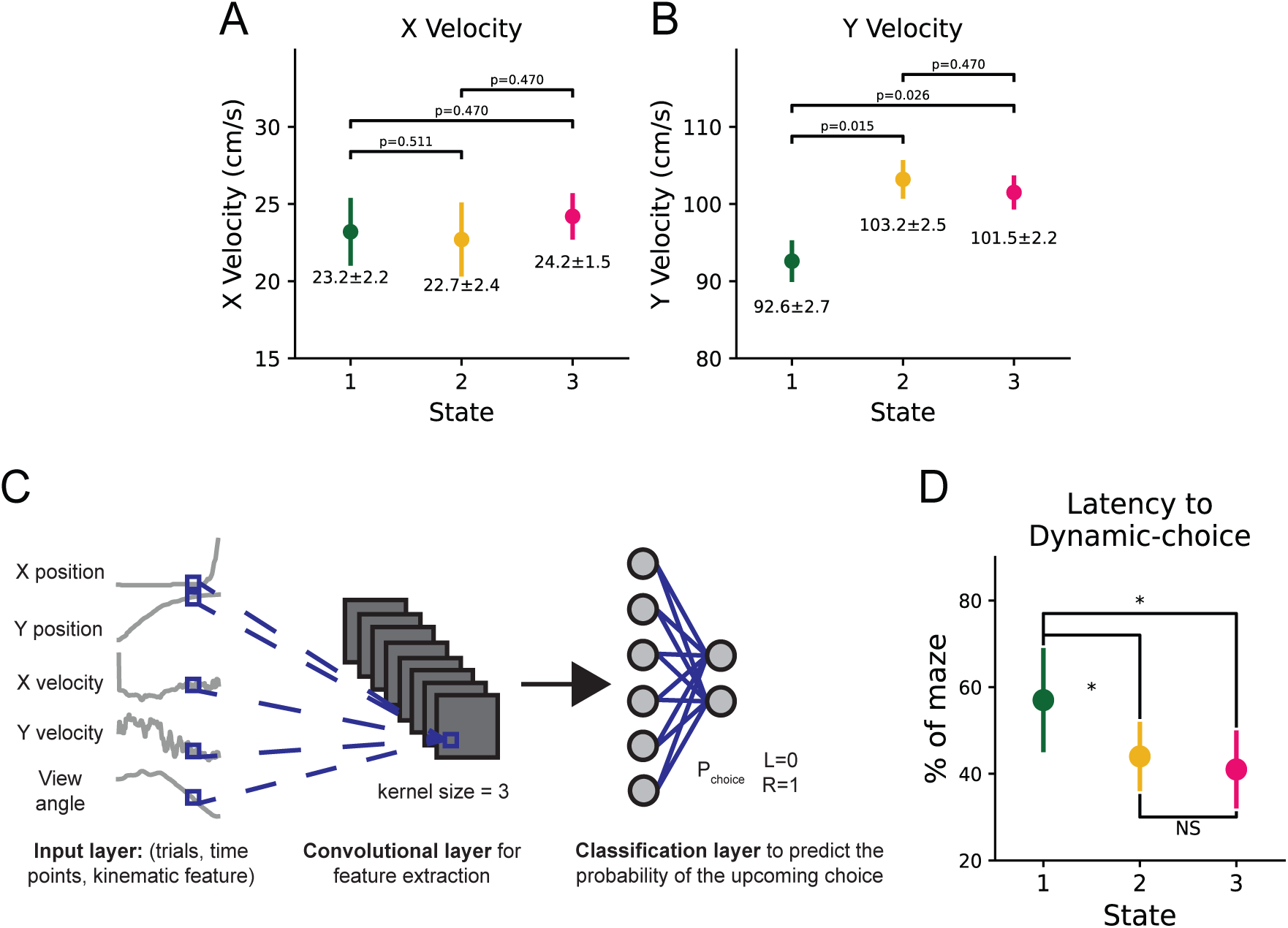
Choice formation was slower in the high-performance state. **(A)** Mean running velocity in the x-axis of the T-maze across the three performance states. **(B)** Mean running velocity in the y-axis of the T-maze across the three performance states. Y-velocity was significantly lower in performance state 1 compared to states 2 and 3 (Mann-Whitney U-test). **(C)** We quantified how well running trajectories in a single trial predicted the mouse’s upcoming choice on that trial using a convolutional neural network (CNN). For each time point in the trial the CNN used running variables such as X and Y position, X and Y velocity, and view angle in the maze from all previous time points to estimate the probability that the mouse turned left or right. **(D)** We then measured the latency to dynamic choice as the time point where the model’s prediction exceeded a threshold of 0.9 for left choice trials and 0.1 for right choice trials. We examined the latency to dynamic choice for all three GLM-HMM latent states across all behavioral sessions and found that the latency to dynamic choice, was significantly higher in the optimal state (state 1) than in the two suboptimal states (state 1 vs state 2, p=0.024. state 2 vs state 2, p=0.011. state 2 vs state 3, p=.680. Mann-Whitney U-test).

**Figure S7.**
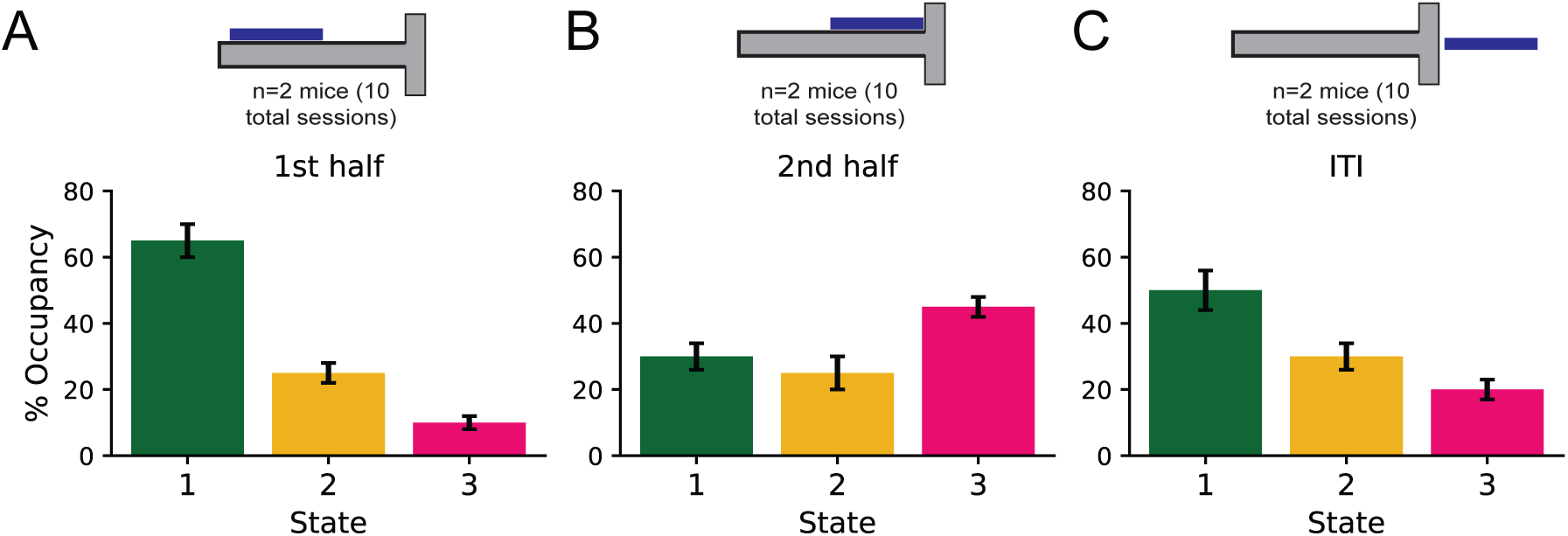
Epoch-based inactivation effects on performance state occupation. **(A)** Fractional occupancies for each state across all behavioral trials during first half inactivation (state 1 = 65.1 ± 5.2, state 2 = 24.6 ± 3.8, state 3 = 11.1 ± 2.6). **(B)** Fractional occupancies for each state across all behavioral trials during second half inactivation. Second half inactivation led to low state 1 occupation and high state 3 occupation. (state 1 = 30.7 ± 4.2, state 2 = 25.3 ± 6.8, state 3 = 45.1 ± 3.9). **(C)** Fractional occupancies for each state across all behavioral trials during ITI inactivation (state 1 = 50.4 ± 6.2, state 2 = 29.7 ± 3.8, state 3 = 21.8 ± 2.2).

**Figure S8.**
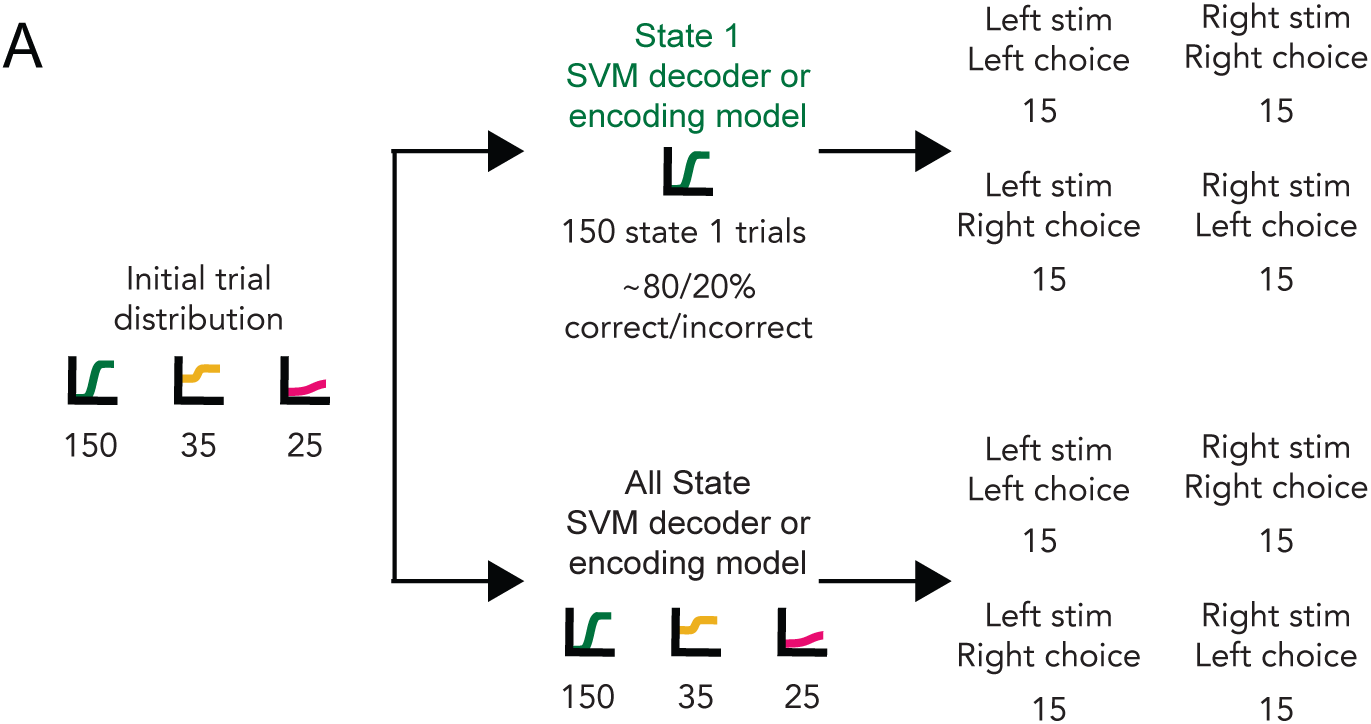
Balancing scheme for neural data analyses. **(A)** Schematic of trial type distribution for all encoding and decoding model (SVM decoders + GLM encoding models). Equal number of left stimulus / left choice, right stimulus / right choice, left stimulus / right choice, right stimulus / left choice trials are used for model training and testing.

**Figure S9.**
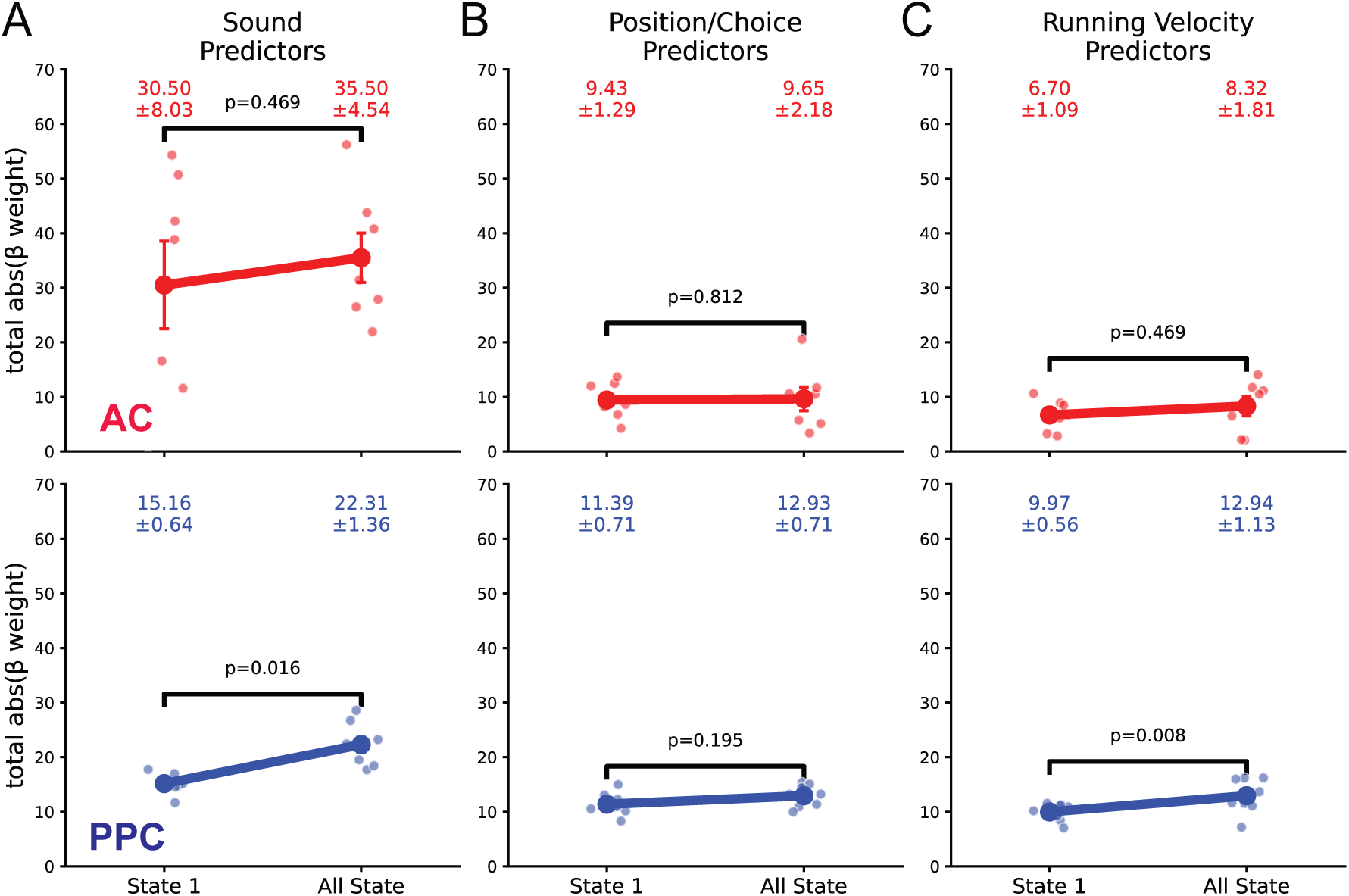
B-weights for the behavior only encoding models for AC and PPC datasets. **(A)** Fitted model weights for variables related to sound location between the two model types averaged across AC (red) and PPC (blue) datasets. (AC: 7 datasets, 329 neurons; PPC: 8 datasets, 4254 neurons, Mann-Whitney U-test). **(B)** Fitted model weights for variables related to position and choice between the two model types averaged across AC and PPC datasets. **(C)** Fitted model weights for variables related to running between the two model types averaged across AC and PPC datasets.

**Figure S10.**
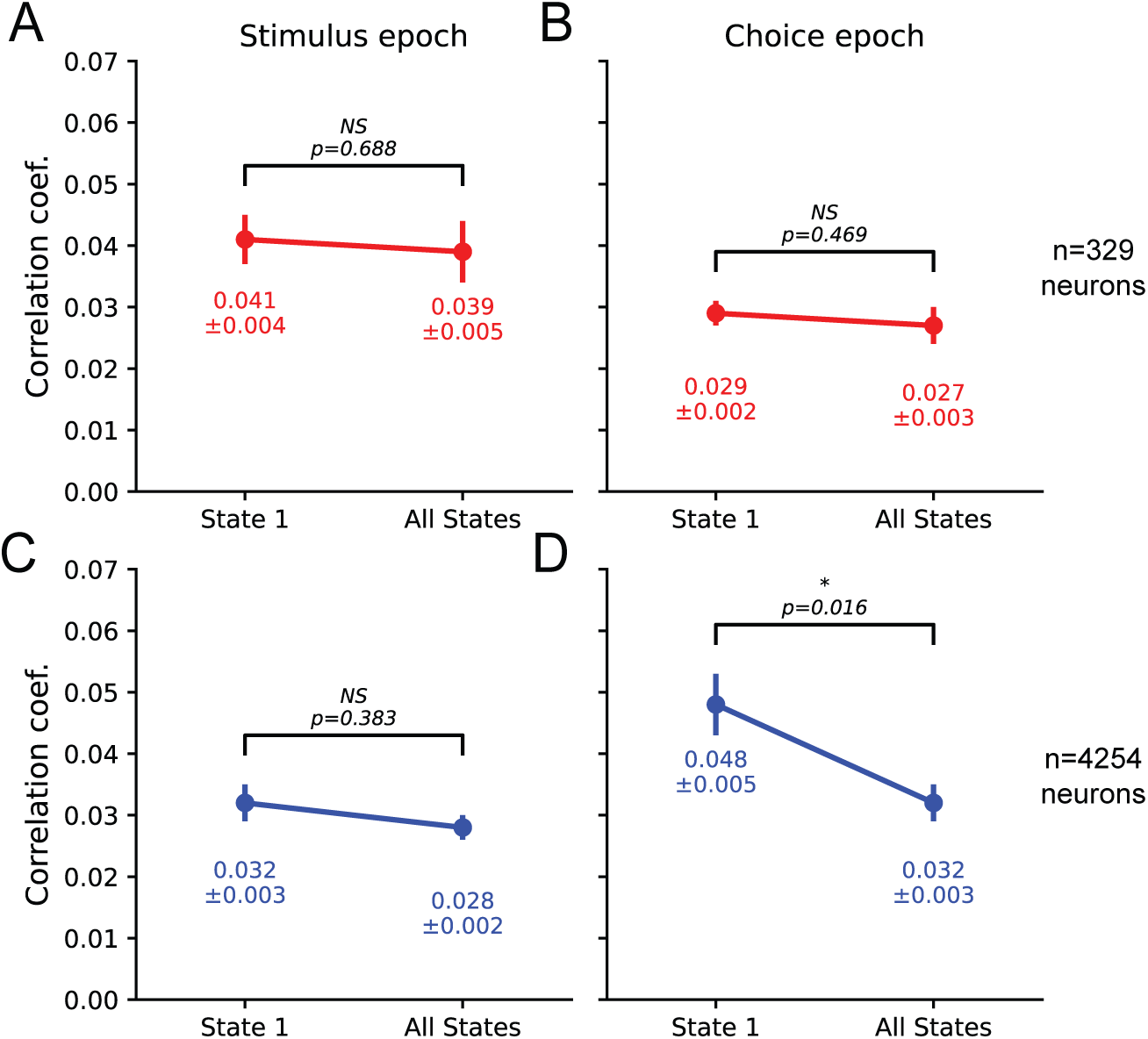
Pairwise noise correlations in AC and PPC neurons during the stimulus and choice epochs. **(A)** Comparison of the pairwise noise correlations between all AC neurons from exclusively state 1 trials or trials spanning all three states during the stimulus epoch (7 datasets, 329 neurons; Mann-Whitney U-test). **(B)** Comparison of the pairwise noise correlations between all AC neurons from exclusively state 1 trials or trials spanning all three states during the choice epoch (7 datasets, 329 neurons; Mann-Whitney U-test). **(C)**Comparison of the pairwise noise correlations between all PPC neurons from exclusively state 1 trials or trials spanning all three states during the stimulus epoch (8 datasets, 4254 neurons; Mann-Whitney U-test). **(D)** Comparison of the pairwise noise correlations between all PPC neurons from exclusively state 1 trials or trials spanning all three states during the choice epoch (8 datasets, 4254 neurons; Mann-Whitney U-test).

**Figure S11.**
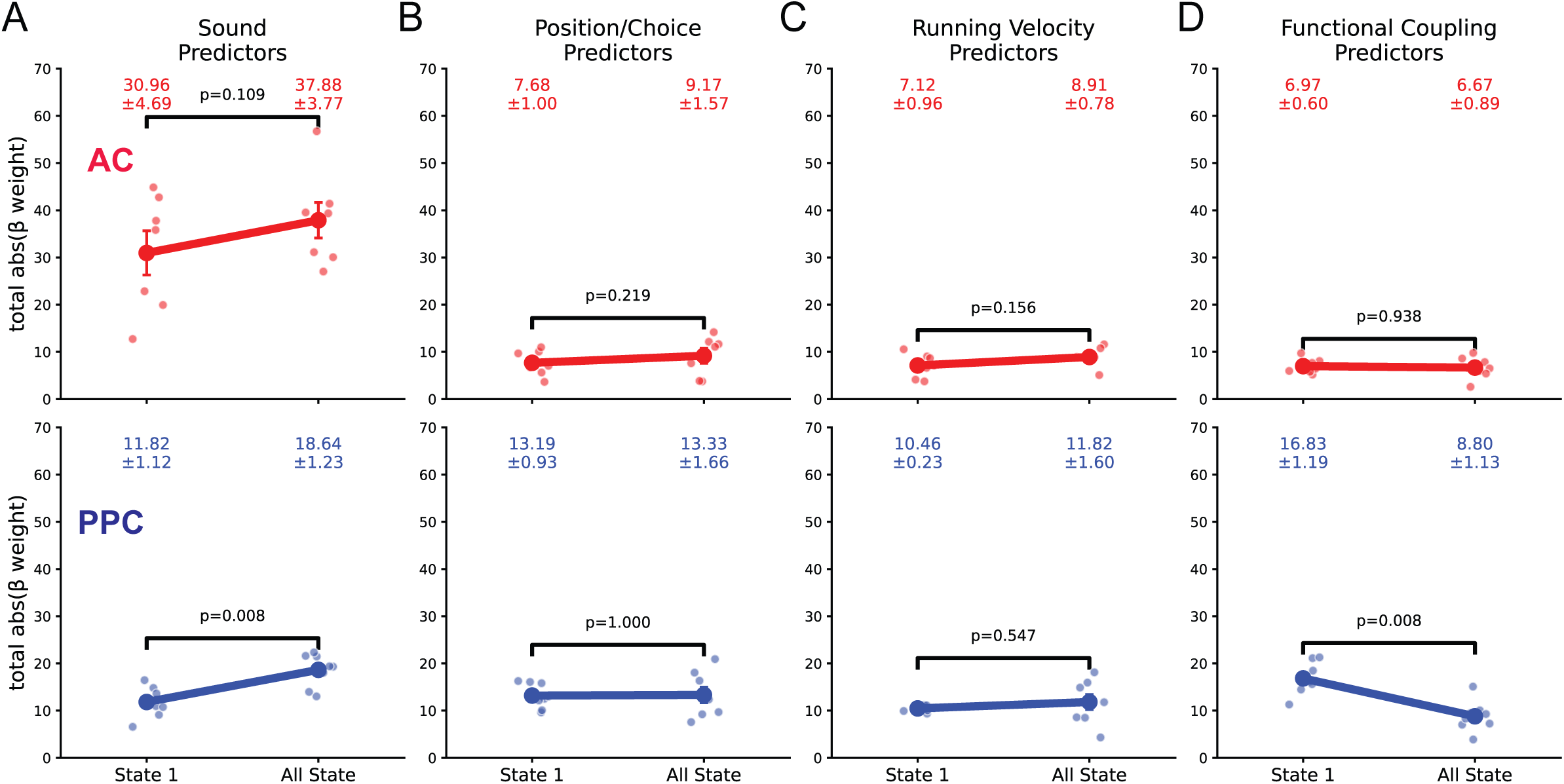
B-weights for the behavior + functional coupling encoding models for PPC and AC datasets. **(A)** Fitted model weights for variables related to sound location between the two model types averaged across AC (red) and PPC (blue) datasets. (AC: 7 datasets, 329 neurons; PPC: 8 datasets, 4254 neurons, Mann-Whitney U-test). **(B)** Fitted model weights for variables related to position and choice between the two model types averaged across AC and PPC datasets. **(C)** Fitted model weights for variables related to running between the two model types averaged across AC and PPC datasets. **(D)** Fitted model weights for functional coupling predictors between the two model types averaged across AC and PPC datasets.

